# Development of an Adaptive, Economical, and Easy-to-Use SP3-TMT Automated Sample Preparation Workflow for Quantitative Proteomics

**DOI:** 10.1101/2025.02.23.639731

**Authors:** Jake N. Hermanson, Lea A. Barny, Lars Plate

## Abstract

Liquid handling robots have been developed to automate various steps of the bottom-up proteomics workflow, however, protocols for the generation of isobarically labeled peptides remain limited. Existing methods often require costly specialty devices and are constrained by fixed workflows. To address this, we developed a cost-effective, flexible, automated sample preparation protocol for TMT-labeled peptides using the Biomek i5 liquid handler. Our approach leverages Single-Pot Solid-Phase-Enhanced Sample Preparation (SP3) with paramagnetic beads to streamline protein cleanup and digestion. The protocol also allows for adjustment of trypsin concentration and peptide-to-TMT ratio to increase throughput and reduce costs, respectively. We compared our automated and manual 18-plex TMT-Pro labeling workflows by monitoring select protein markers of the Unfolded Protein Response (UPR) in pharmacologically activatable, engineered cell lines. Overall, the automated protocol demonstrated equivalent performance in peptide and protein identifications, digestion and labeling efficiency, and an enhancement in the dynamic range of TMT quantifications. Compared to the manual method, the Biomek protocol significantly reduces hands-on time and minimizes sample handling errors. The 96-well format additionally allows for the number of TMT reactions to be scaled up quickly without a significant increase in user interaction. Our optimized automated workflow enhances throughput, reproducibility, and cost-effectiveness, making it a valuable tool for high-throughput proteomics studies.

## Introduction

Advancements in mass spectrometry (MS) instrumentation has enabled the study of proteins with high sensitivity and reproducibility in both basic and translational research at an unprecedented scale^1,2^. Regardless of the application, reproducible sample preparation is essential for minimizing technical variability to avoid obscuring biological insight^3^. As MS analysis speed increases, sample preparation has become a bottleneck for throughput. To address this challenge, liquid handling robots have been developed by various vendors: Agilent^4^, ThermoFisher^5^, Beckman Coulter^6–8^, and others to automate key steps in the bottom-up proteomics workflow—including reduction, alkylation, cleanup, digestion, and acidification, following initial user interaction^9^. While numerous automated liquid handling protocols have been developed for label free peptide generation, few protocols have been developed for the generation of isobarically labeled peptides^7,10^. Isobaric labeling using commercial tandem mass tags (TMT) allows for multiplexing of up to 35 samples for concurrent measurement in a single MS analysis, significantly increasing analysis throughput^11–13^.

Previous methods for the preparation of TMT-labeled peptide samples on automated liquid handling systems have required specialty devices such as a vacuum manifold and positive-pressure apparatus to automate solid phase extraction based cleanup steps^7^. However, these additions result in higher overall instrument cost and may not be widely applicable to other automated protocols utilized by the laboratory. In contrast, instruments like the Accelerome (ThermoFisher Scientific) have been specifically developed for label-free and isobaric-tagged bottom-up proteomics sample preparations. However, this instrument can only perform preprogrammed workflows and the protocol requires purchasing a commercial kit limited to the preparation of 36 samples per cycle^10^.

To address the need for a flexible and cost-effective automated sample preparation method for TMT labeled peptides, we developed an automated workflow on a Biomek i5 liquid handler (Beckman Coulter). Our method begins with unnormalized cell lysate and produces either 16-or 18-plex TMT-labeled samples ready for MS analysis (**Fig 1A**). For protein cleanup and digestion, our protocol uses Single-Pot Solid-Phase-Enhanced Sample Preparation (SP3).^14^ The protocol leverages commercially available SP3 beads with specialized surface chemistries for protein binding^15^. The paramagnetic property of these beads make them particularly conducive to automation and wash steps needed to remove detergents, buffers, etc^4^. Additionally, our automation protocol allows for the adjustment of trypsin concentration to shorten the digestion incubation^16^ and variation of peptide-to-TMT concentration to increase the number of labeling experiments that can be performed with a set of purchased TMT reagents^17^. The Biomek i5 may serve as an attractive liquid handling robot for labs as it can also be adapted to perform numerous other tasks, including: cell culture^18^, RNA/ DNA extractions^19^, and versatile assay development. Utilizing our 96-well automation format in combination with TMT isobaric tags enables the parallel preparation of six 16-plex or four 18-plex multiplexed MS pools at a rate of 10.8 mins per sample (TMT channel) which is less than the Accelerome at 12.6 mins per sample (**Fig 1B, Supplemental Table 1**).

**Figure 1.**
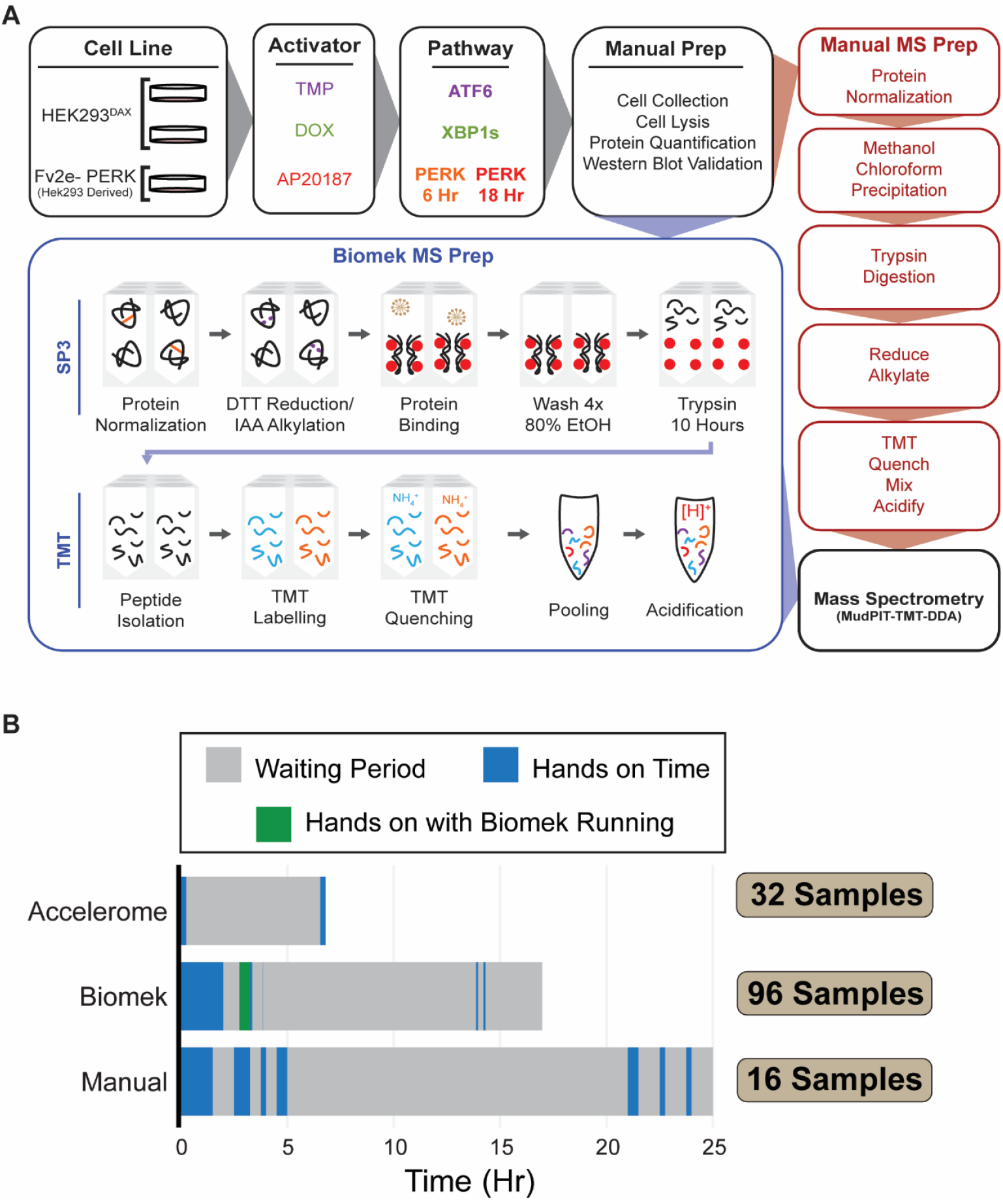
Overview of the Biomek i5 and manual TMT-labeling methods and comparison of the time required to carry out the protocols discussed. **(A)** Outline of the Biomek and manual preparation method used in this study for the generation of TMT-labeled peptides. The Biomek automated method used SP3 beads for protein clean-up and digestion, while the manual protocol employed a Methanol/Chloroform precipitation technique. (**B**) Comparison of the time (hr) required to prepare TMT labeled peptides using either the Accelerome (ThermoFisher Scientific), Biomek i5 (Beckman Coulter), or our manual method. Hands-on time includes preparing reagents (both on Biomek and manually) and, for the Biomek specifically, loading samples onto the deck and setting up pipette tips.

We validated our automated workflow by determining activation of the Unfolded Protein Response (UPR), a biologically important pathway with implications in diabetes^20,21^, neurodegeneration^22–24^, and cancer^25–27^. The UPR is triggered by the accumulation of misfolded and aggregated proteins in the endoplasmic reticulum (ER). This response involves three distinct pathways: PERK, ATF6, and IRE1/XBP1s (**Fig 2**)^28–31^. Activation of each branch of the UPR results in a distinct transcriptional and translational response^31–34^ in which pathway-specific genes and proteins are subsequently upregulated. Branch-specific protein markers can be monitored via liquid chromatography-MS/MS (LC-MS/MS) to define the state of the repsonse^35,36^. To elicit the UPR response branch-specifically, engineered stable cell lines which can be activated with small molecules were utilized^32^. We then prepared TMT-Pro labeled 18-plex samples using the automated pipeline and a conventional manual protocol^37–39^. The manual and automated workflow showed a similar number of peptides and proteins identified with equivalent precision across treatments. Additionally, similar tryptic digestion, alkylation, and TMT labeling efficiencies were observed between the methods. The Biomek preparation method resulted in higher abundances of many core UPR target proteins, highlighting an enhanced quantification dynamic range, while the coefficient of variation (CV) for these proteins remained consistent between the methods. Lastly, compared to the manual TMT label preparation, our automated workflow significantly decreased hands-on time from 24 to 2.8 hours, respectively, for the generation of 96 samples. The reduction in hands-on time minimizes the risk of sample mix-ups and reagent addition errors. Implementing our developed workflow will enhance the throughput of samples processed for MS analysis in a reproducible and cost-effective manner.

**Figure 2.**
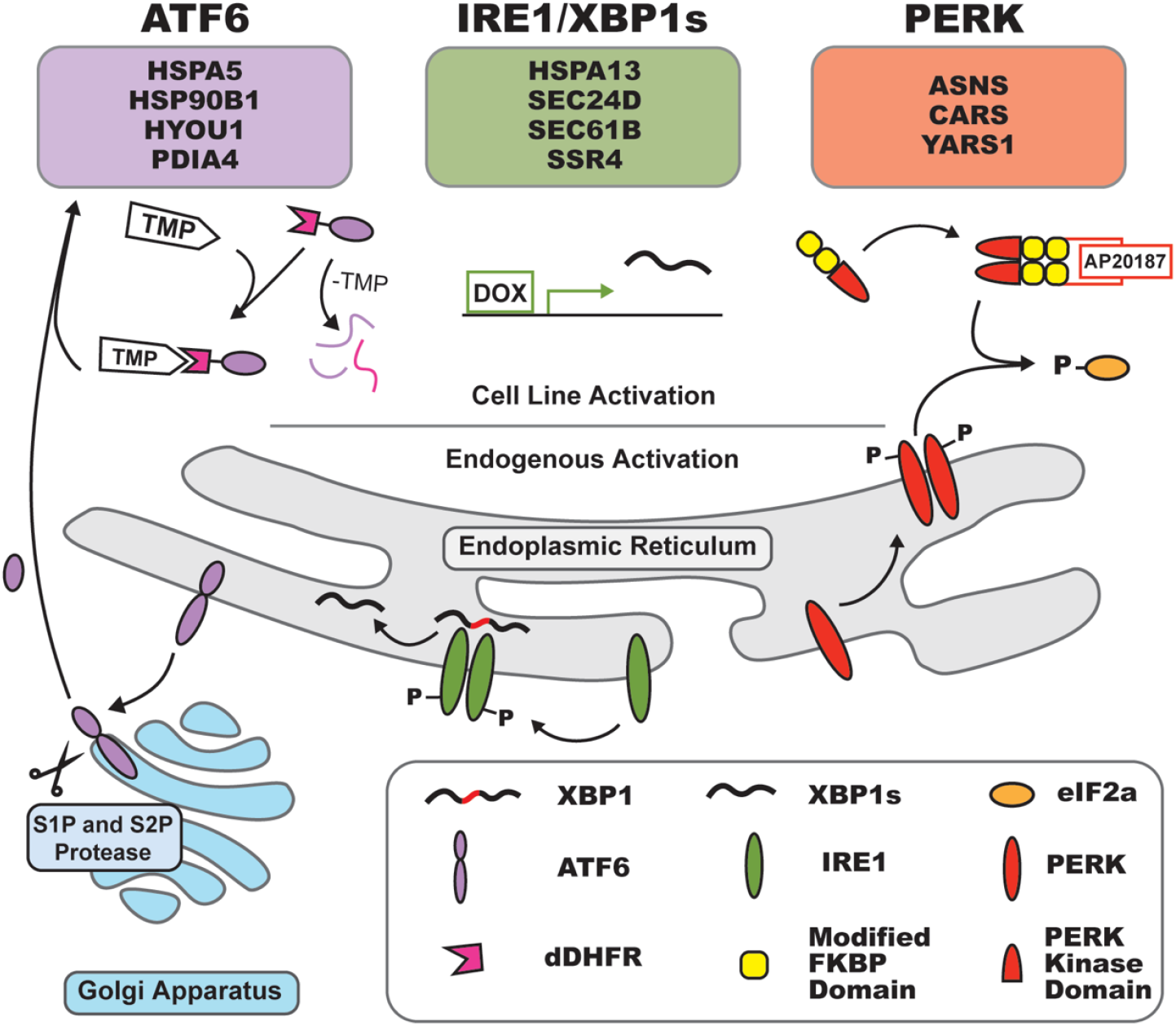
Overview of the three branches of the Unfolded Protein Response (UPR). Known branch-specific UPR target protein markers were used throughout this study to monitor the state of the response. At the top of the figure, several genes are highlighted that are upregulated during activation. ATF6 activation causes upregulation of HSPA5, HSP90B1, HYOU1 and PDIA4. HSPA13, SEC24D, Sec61B and SSR4 are upregulated by XBP1s, while ASNS, CARS, and YARS1 are upregulated upon PERK activation. Endogenous activation of ATF6 occurs when ATF6 translocates from the endoplasmic reticulum to the golgi apparatus. Cleavage by S1P and S2P proteases occur here to release ATF6-N. This truncated form of ATF6 then moves to the nucleus and acts as a transcription factor. This endogenous activation is circumvented by tagging the active ATF6-N with destabilized DHFR (dDHFR). This tag targets ATF6-N for degradation, but when treated with TMP stabilizes dDHFR allowing for ATF6-N to act as a transcription factor. IRE1/XBP1s is activated when IRE1 autophosphorylates and non-canonically splices the XBP1 mRNA into XBP1s which is translated into a transcription factor. In the HEK^293^DAX engineered stable cell line, a doxycycline (dox)-inducible Tet-On system is used where XBP1s is transcribed in the nucleus and does not require additional splicing. Lastly, endogenous PERK becomes active when it dimerizes, autophosphorylates and in turn phosphorylates eIF2a. Activation of the Fv2e-PERK cell line occurs with a fusion protein containing the PERK kinase domain and two FKBP domains that when treated with AP20187 cause dimerization and subsequent activation. The ATF6, XBP1s, and PERK branches of the UPR were pharmacology activated with TMP (10 μM), DOX (1 μg/mL), and AP20187 (5 nM) for 16-18 hours, respectively.

## Results and Discussion

### Establishing an Instrument Workflow on the Biomek i5 for the Generation of TMT-Labeled Peptides

The Biomek i5 is a versatile liquid handler capable of automating sample preparation for a variety of applications, including genomic analysis (qPCR, transcriptomics, etc)^19,40^ and LC-MS/MS workflows, particularly for proteomics.^7,8,41,42^ Here, we present an optimized protocol using this instrument to prepare 96 TMT labeled peptide samples ready for mass spectrometry (**Fig. 1A**). Briefly, the protocol begins with lysate normalization following off-line determination of protein concentration in whole-cell lysates using a bicinchoninic acid assay.^43^ A CSV file containing the buffer and lysate amounts for normalization to 20 μg of protein is then uploaded to the instrument for automated normalization into a deep-96 well plate. Proteins are then reduced and alkylated using dithiothreitol (DTT) and iodoacetamide (IAA), respectively, and subsequently bound to SP3 paramagnetic beads for cleanup prior to digestion with trypsin/Lys-C. For steps requiring temperature control and/ or shaking, the Bioshake Q1 Heater Cooler Shaker (QINSTRUMENTS) was used. For example, during normalization and after protein digestion samples were held at 4°C until further user interaction. Samples can then either be collected for label-free mass spectrometry or remain on the Biomek i5 for TMT labeling, quenching, pooling, and acidification.

In contrast, the manual preparation involved normalizing samples by hand and performing methanol/chloroform precipitation for protein isolation. Briefly, proteins were resolubilized in 1% RapiGest and reduced with tris(2-carboxyethyl)phosphine (TCEP), followed by alkylation with IAA. All other steps in the TMT-labeled peptide generation process mirrored the Biomek i5 protocol but were executed manually instead. Initial iterations of this protocol involved the movement of RapiGest resuspended proteins onto the Biomek following methanol/chloroform precipitation for subsequent reduction, alkylation, digestion and TMT labeling. While this method resulted in high peptide and protein identifications, transfer of protein in this manner was extremely time consuming and risked protein loss, prompting the need to optimize an automated SP3 method. SP3 beads have the added benefit of being compatible with low protein inputs.^44^ Following digestion, paramagnetic SP3 beads are easily precipitated from peptides using a removable universal magnet plate without the need for further peptide clean-up steps.

Following digestion, TMT-Pro labeling of peptides was carried out either manually or on the Biomek. To facilitate easier storage and dispensing on the Biomek, the TMT reagents were stored in LVL tubes. LVL tubes are individual, barcoded tubes with screw caps that are available for purchase in a 96-well format. These tubes can be filled with variable quantities of TMT reagent, providing flexibility in reagent storage and dispensing. While this method is convenient, it is optional. Alternatively, TMT reagents can be manually transferred to a 96-deep-well plate or purchased pre-packaged in a 96-well format. Our instrument is additionally outfitted with a High Efficiency Particulate Air (HEPA) filtration system to prevent sample contamination from air particles. Overall, the only specialized devices required to run our protocol are the Bioshake Q1 Heater Cooler Shaker and removable magnetic plate.

One key difference between the methods described is the time required to generate a single TMT-labeled sample. The entire Biomek and manual protocol generates TMT labeled samples at a rate of 10.8 and 93.6 mins/ sample, respectively (**Fig. 1B**). In addition to passive incubations in which the user is not required to engage with the instrument or method, indicated as a “waiting period” in Figure 1B in gray, the Biomek protocol also requires manual reagent preparation (bead washing, trypsin, etc) while the instrument is performing reduction and alkylation steps, indicated in green as “hands on with Biomek running” (**Fig. 1B**). The ability to add reagent to all wells in use on the Biomek significantly decreases the manual time required for sample preparation. Overall, the Biomek-assisted protocol requires approximately 2.8 hours of manual intervention to generate 96 TMT-labeled samples, compared to 24 hours for the fully manual protocol. Alternatively, the Accelerome (ThermoFisher Scientific) generates samples at a rate of 12.6 mins/ sample. The sample production rate is reduced using the Accelerome compared to the Biomek largely due to the number of samples that can be prepared at a given time (**Fig. 1B**).

### Comparing the Performance of the Preparation Techniques: Manual vs. Biomek i5

To validate our developed Biomek protocol, we utilized established engineered stable cell lines (HEK293^DAX^ and Fv2e-PERK) in which the three branches of the UPR can be activated individually with small molecules (trimethoprim (TMP), doxycycline (DOX), AP20187 (cell-permeable dimerizer of FK506-Binding Protein (FKBP) fused proteins)), allowing us to test the differences between the manual and Biomek i5 preparation protocols (**Fig. 2**)^32,45^. We employed well-characterized antibody-based markers for branch-specific UPR activation, such as BiP for ATF6 activation and eIF2α-P for early PERK activation (6 hours), followed by ASNS upregulation for late PERK activation (18 hours). UPR activation in cell lines was confirmed by western blotting prior to MS analysis (**Supplemental Fig. 1**)

Both manual and Biomek prepared TMT 18-plex samples were analyzed using Multidimensional Protein Identification Technology (MudPIT) – data dependent acquisition (DDA)^46^. MudPIT utilizes ion exchange chromatography to fractionate complex peptide mixtures, enhancing peptide detection^47–49^. For each analysis, identical amounts of peptides were loaded on-column. To determine the number of peptide groups and proteins identified using the two sample preparation protocols, we first removed known common contaminants^50^ and retained proteins with a minimum of two unique peptides. Using these criteria, 24,061 peptide groups were identified with manual preparation, while 24,052 peptide groups were identified with the Biomek i5 preparation (**Fig. 3A**). Alternatively, 3,544 and 3,676 proteins were identified using the manual and Biomek i5 automated preparation, respectively (**Fig. 3B, Supplemental Table 2 & 3**). High overlap was observed between the methods in identified peptide groups and proteins (**Fig. 3C/D**).

**Figure 3.**
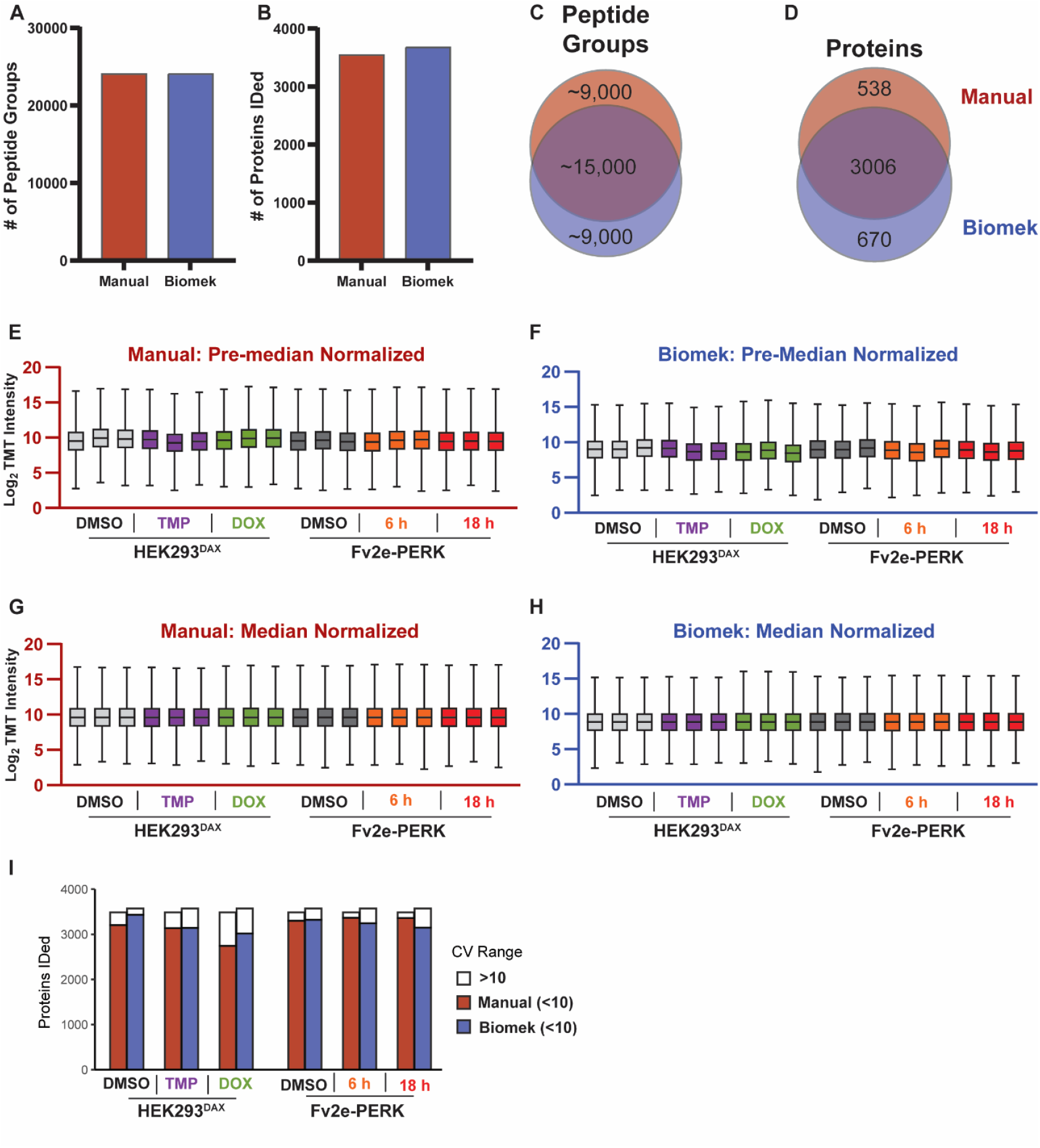
Biomek and manual sample preparation show similar performance in peptide and protein identifications. **(A/B)** Bar graphs indicate the number of peptide groups and proteins detected, in all TMT channels, using either the manual or Biomek i5 preparation. **(C/D)** Unique and shared peptide groups and proteins identified between the two methods. Distribution of protein abundances pre **(E/F)** and post median normalization **(G/H).** Coefficient of variation (CV %) ranges of the proteins identified in each of the cell lines with activation, comparing the TMT sample preparation methods.

To compare the precision of the workflows, we assessed the median abundances of each sample before and after median normalization, along with the distribution of protein CVs. When comparing the distribution of protein abundances prior to normalization, neither preparation had samples that deviated far from the average median, indicating the methods did not result in high sample loss (**Fig. 3E/F**). After median normalization, all samples had a similar median abundance, regardless of the sample preparation technique utilized (**Fig. 3G/H**). Lastly, CVs were calculated for all identified proteins on the non-log-transformed intensities of biological replicates, grouped by treatment and cell line, after applying global median normalization using the equation CV= σ (mean)/ μ (standard deviation).^51^ The number of proteins with a CV less than 10 (indicating high precision) was similar between the methods, however the Biomek preparation had only slightly more proteins that were identified with CVs greater than 10 (9.9% vs. 8.6%) (**Fig. 3I**). Overall, these methods show similar precision in their preparations.

### Comparing Efficiency of Trypsin Digestion, Alkylation, and TMT Labeling

To better understand the potential differences in sample preparation between the manual and automated protocols, we assessed the efficiencies of digestion, alkylation, and TMT labeling. First, to evaluate trypsin digestion efficiency, we compared the number of missed cleavages between the two TMT peptide preparation protocols. Both protocols resulted in a similar number of missed cleavages, indicating that the digestion time and trypsin concentration utilized was suitable for the trypsin-to-lysate ratio (**Fig. 4A**). In our protocol, trypsin/lys-C was utilized at a 1:40 enzyme/ substrate ratio for 10 hours. Importantly, higher enzyme concentrations can be utilized to reduce digestion times, allowing for an additional gain in sample preparation throughput.^52,53^ This approach, however, is more costly compared to the enzyme concentrations used in this study.

**Figure 4.**
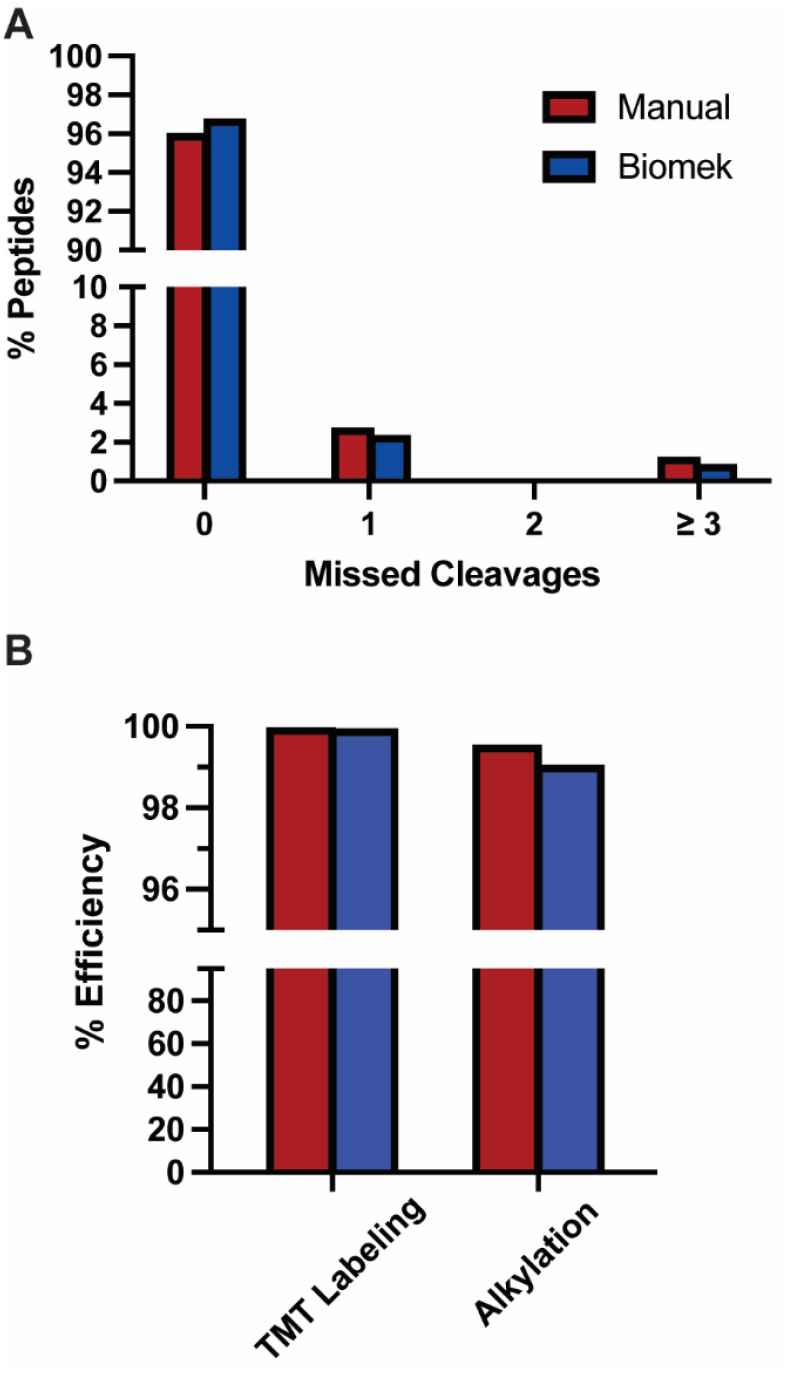
Biomek and manual preparation show comparable labelling and digestion efficiencies. **(A)** The percentage of peptides with zero, one, two, or more than three trypsin/lys-C missed cleavages detected in both manual and Biomek i5 TMT sample preparations. **(B)** TMT labeling and alkylation efficiency of the two preparations.

Alkylation and TMT labeling efficiencies were evaluated by searching the data with carboxyamidomethylation and TMT-Pro modifications, respectively, as dynamic rather than static variables to determine the number of peptides missing the desired modification. Importantly, the methods did not result in different alkylation or TMT labeling efficiencies (**Fig. 4B**). The methods differed minimally in alkylation efficiency by 0.75%. Additionally, both preparations yielded TMT labeling efficiencies over 99%. These data indicate that the quality of sample preparation is consistent, whether performed manually or using the automated protocol. Importantly, our TMT labeling method differs from the manufacturer’s protocol in that the peptide to TMT reagent ratio was 1:5 as opposed to the recommended 1:25. Previously, it was reported that peptide-to-TMT ratios as low as 1:1 (weight/weight) result in labeling efficiencies exceeding 99%.^17^ Our automated TMT labeling approach makes the technique cost-effective and scalable for large-scale studies.

### Evaluating TMT-Based Quantification in the Context of the Unfolded Protein Response (UPR)

To determine whether the automated or manual preparation provided better quantification accuracy the upregulation of known protein markers for each branch of the UPR were assessed^32,35^. For the majority of branch specific UPR markers evaluated, the log_2_ fold change of the treatment compared to the DMSO control was greater for the Biomek preparation than that of the manual preparation (**Fig. 5A, Supplemental Fig. 2A/B**). This was especially pronounced for the ATF6 branch of the UPR pathway. We thus compared ATF6 activated samples prepared either manually or with the Biomek to reveal higher abundances for many of the prominent upregulated ATF6 markers in the Biomek preparation (**Fig. 5B**). These data suggest that the method of protein cleanup and isolation (e.g., SP3 or Methanol/Chloroform) can influence the recovery of specific proteins, especially those with a high dynamic range, such as those upregulated during the UPR response.^54^ The ATF6-associated proteins, namely HSPA5, HSP90B1, and PDIA4, exhibited the greatest differences in abundance between the two preparation methods in TMP-treated HEK293^DAX^ cells, and were also the most differentially expressed proteins in the dataset. Similar trends were observed for proteins regulated by XBP1s and PERK (assessed at 6 h and 18h), although the differences in abundance between the two methods were less pronounced, likely because these proteins are not as strongly differentially expressed (**Supplemental Fig. 2C/D**).

**Figure 5.**
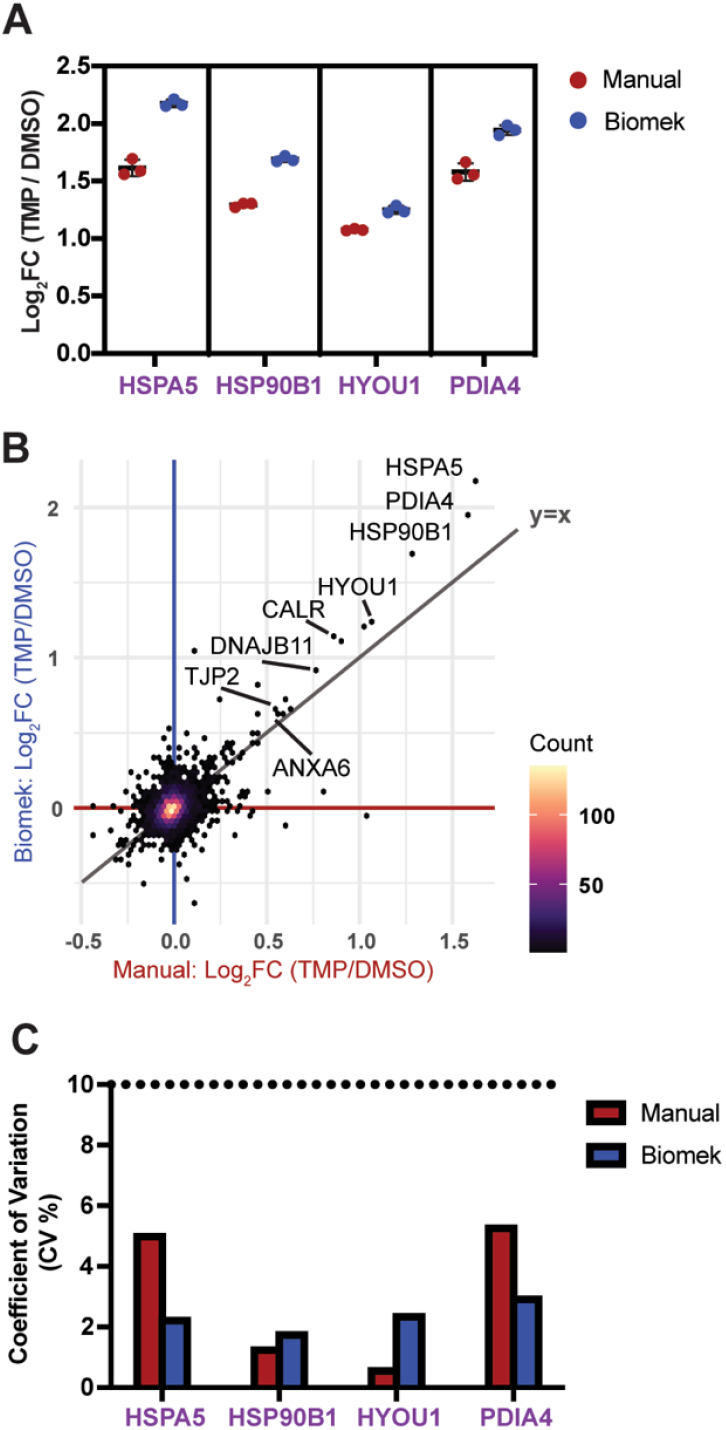
Identification of core UPR protein markers in the Biomek and manual preparations. **(A)** Log_2_ fold change of core ATF6 target proteins when prepared with the two methods. **(B)** Correlation plot of the log_2_ fold change of TMP activated HEK^293^DAX cells compared to DMSO in both preparation methods. (**C**) Coefficient of variation of select AFT6 regulated UPR target proteins.

Secondly, the CV of the branch-specific UPR proteins was monitored to assess whether the methods differed in the variation introduced during sample preparation **(Fig. 5C)**. Low variability (CV% < 10) was observed within biological replicates of the sample condition, indicating that the methods provide robust sample preparation (**Supplemental Fig. 2E/F**). Overall, these data reveal the SP3 beads preparation, executed on the Biomek, promotes identification of select UPR markers with high precision. As these markers in different biological contexts may be upregulated to different extents, a technique that is able to accommodate a large dynamic range is preferable.

## Conclusion

Here we introduce an automated workflow designed to expedite the acquisition of TMT based proteomics data. This process converts unnormalized cell lysates into reduced and alkylated TMT-labeled peptides with minimal user intervention, utilizing SP3 paramagnetic beads. While users must prepare reagents for loading onto the Biomek, the hands-on time is significantly reduced and remains consistent for up to 96 samples. The Biomek-assisted protocol requires approximately 2.8 hours of manual intervention to generate 96 TMT-labeled samples, compared to 24 hours for the fully manual protocol. The ability to process 96 samples in tandem on the Biomek provides considerable time savings over existing methods on other commercial platforms. Additionally, we demonstrate that the data quality achieved with our automated pipeline is improved by increasing the dynamic range of TMT quantification changes. This approach enhances data acquisition efficiency without necessitating extensive hands-on training or experience.

## Materials and Methods

### Cell Culture and Harvesting

HEK293^DAX^ and HEK293 Fv2e-PERK cells were previously established as engineered stable cell lines to study the UPR and were a kind gift from Luke Wiseman (Scripps Research Institute)^32,45^. Cells were cultured in Dulbecco’s modified Eagle medium supplemented with 10% fetal bovine serum, 1% penicillin/streptomycin and 1% glutamine (Incubated at 37 °C with 5% CO_2_). To induce the upregulation of ATF6 or XBP1s transcription factors in the HEK293^DAX^ cell line, cells were incubated with either 10 μM TMP or 1 μg/mL doxycycline (DOX), respectively, for 18 hours. To promote PERK transmembrane receptor dimerization, Fv2e-PERK cells were treated with 5 nM of AP20187 and were harvested at both 6 and 18 hours post drug addition. Cells were washed 2x with ice cold phosphate buffered saline (PBS), then scraped and collected in ice cold PBS + 1mM ethylenediaminetetraacetic acid (EDTA), followed by cell pelleting (200 g, 5mins). The pellet was washed 2x with cold PBS and centrifuged at 200 g for 5 min. Remaining PBS was removed by suctioning followed by addition of radioimmunoprecipitation assay (RIPA) buffer (50 mM Tris (pH 7.5), 150 mM NaCl, 0.1% SDS, 1% Triton X-100, 0.5% deoxycholate) containing 1x Protease inhibitor (Roche complete EDTA-free protease inhibitor Millipore Sigma, 4693132001; HEK293^DAX^ cells) and 1x Protease + 0.1% (v/v) phosphatase inhibitor (Fv2e-PERK cells) for 15 min. Insoluble debris was then pelleted via centrifugation (16 kg, 15 mins) followed by quantification of protein levels via bicinchoninic acid assay^43^.

### Automated Sample Preparation

The method for the Biomek i5 system can be found in the supplement. Cell lysate was normalized to 20 μg of protein in 70 μL RIPA buffer on the Biomek i5 followed by SP3 bead digestion and cleanup described elsewhere^14^. Briefly, protein was digested on 8 μl total of SP3 beads (1:1 hydrophilic to hydrophobic beads) in 50 μL of 50 mM HEPES (pH 8) with Trypsin/Lys-C in a 1:40 protease to protein ratio for 10 hours at 37 °C/ 700 rpm. After digestion, peptides were held at 4 °C until TMTPro labeling. To begin the labeling protocol, peptides were first transferred to a new PCR plate (free of SP3 beads) followed by the addition of 10 μL H_2_O. Peptides were then reacted with 40 μL of resuspended TMTPro 18-plex reagent for 1 hour at room temperature. Purchased TMTPro reagent was dissolved in acetonitrile (ACN) to achieve a final concentration of 2.42 mg/mL. For ease of use on the automated protocol, TMT reagents were stored in MX500 LVL tubes (Thomas Scientific). The TMT reaction was subsequently quenched with the addition of fresh 10% ammonium bicarbonate (w/v in H_2_O) to a final concentration of 4% (v/v) and incubated for 1 hour at room temperature. Samples were then pooled into a fresh LoBind tube (ThermoFisher Scientific) and acidified using formic acid (FA, ThermoFisher Scientific, PI28905) to ensure pH ≤ 2.0. The pooled sample was then dried *in vacuo* using a SpeedVac (Thermo Fisher) until a third of the volume total (after labeled peptide pooling/ acidification) remained. The volume of the sample was then adjusted back to the original pooled volume with buffer A (95% H2O, 4.9% ACN, 0.1% FA), centrifuged for 30 min at 21.1 kg to remove residual beads, and the supernatant was transferred to a fresh LoBind tube prior to LC-MS/MS analysis.

### Manual Sample Preparation

Cell lysate was normalized to 20 μg of protein in 100 μL RIPA buffer followed by methanol/chloroform precipitation. Protein was precipitated by adding methanol (MeOH), chloroform, and water in a 3:1:3 ratio, respectively. The sample was then vortexed and spun down for 1 min at 14k RPM, followed by removal of the top layer and two additional washes with 500 μL MeOH. After the final wash, MeOH was suctioned off and samples were then left to air dry. Protein pellets were then resuspended in 5 uL 1% RapiGest SF (w/v; Waters #186002122) surfactant followed by an addition of 10 uL of 0.5 M HEPES (pH 8.0). Volume was then adjusted to 47.5 uL with H_2_O. Reduction of proteins was carried out with 5 mM TCEP (30 mins at RT) followed by protein alkylation (30 mins, dark at RT) using 10 mM iodoacetamide. Next, proteins were digested overnight (10 hours) at 37 ºC using Trypsin/Lys-C at a 1:40 protease to protein ratio while shaking at 750 rpm. Samples were TMT labeled in the same manner as described in the *Automated Sample Preparation section*. Lastly, samples were pooled and acidified with FA (added to reach pH 2). The sample was then concentrated on the SpeedVac to one-third of the pooled volume to remove ACN. The volume of the sample was then adjusted back to the original pooled volume with buffer A, centrifuged for 30 min at 21.1 kg to remove RapiGest SF, and the supernatant was transferred to a fresh lobind tube prior to LC-MS/MS analysis.

### Liquid Chromatography-Tandem Mass Spectrometry

LC-MS/MS data-dependent analysis was performed using an Exploris480 mass spectrometer (Thermo Fisher) equipped with a Dionex Ultimate 3000 RSLCnano system (Thermo Fisher). MudPIT columns were made as previously described^49^ with 15 μg of labeled peptides loaded onto the trap column using high pressure chamber and washed for 30 mins with Buffer A prior to MS analysis. To elute peptides from the first C18 phase to the SCX resin of the MudPIT column, 10 μL of Buffer A was injected and the following 90 minute gradient was utilized: 2% B (5 min hold) ramped to a mobile phase concentration of 40% B over 35 minutes, ramped to 80% B over 15 mins, held at 80% B for 5 mins, then returned to 2% B in 5 mins and then held at 2% B for the remainder of the analysis at a constant flow rate 500 nl/min. Then to fractionate the sample, 10 μL sequential injections of 10, 20, 30, 40, 50, 60, 70, 80, 90 and 100% buffer C (500 mM ammonium acetate in buffer A) in Buffer A, with a final fraction injection of 90% buffer C and 10% buffer B (99.9% acetonitrile, 0.1% formic acid v/v) were utilized with a corresponding 130 min gradient at a flow rate of 500 nl/min (4% B for 10 mins then ramped to 40% B over 90 mins, increased to 80% B over 5 mins then held at 80% B for 5 mins and returned to 4% in 5 mins, and held at 4% for the remainder of the analysis). For TMT-DDA acquisition, a 3 sec duty cycle was utilized consisting of a full scan (400-1600 *m/z*, 120,000 resolution) and subsequent MS/MS spectra collected in TopSpeed acquisition mode. For MS^1^ scans, the maximum injection time was set to 50 msec with a normalized AGC target of 100%. Ions were selected for MS/MS fragmentation based on the following criteria: MS^1^ intensity above 1e4, charge state between 2-6, and monoisotopic peak determination set to peptide. Additionally, a dynamic exclusion time of 45 secs was utilized (determined from peptide elution profiles) with a mass tolerance of +/-10 ppm to maximize peptide identifications. MS/MS spectra were collected with a normalized HCD collision energy of 36, 0.4 m/z isolation window, auto selected for maximum injection time mode, a normalized AGC target of 100%, at a MS^2^ orbitrap resolution of 45,000 with a defined first mass of 110 *m/z* to ensure measurement of TMTPro reporter ions.

### Data analysis

Proteome Discoverer 2.4 (Thermo Fisher Scientific) was utilized to obtain peptide identifications and TMT-based protein quantification. MS/MS spectra were searched using SEQUEST-HT against a Uniprot SwissProt canonical human FASTA database (downloaded 01May2024; containing 20,361 entries), contaminant FASTA^50^ (containing 379 entries), and a decoy database of reversed peptide sequences. Peptide precursor ion mass tolerance was set to 20 ppm, and fragment ion mass tolerance was fixed at 0.02 Da. Only peptides with a minimum length of six amino acids were considered, and trypsin digestion was assumed with a maximum of two missed cleavages allowed. Dynamic modifications included oxidation of methionine (+15.995 Da), protein N-terminal methionine loss (-131.040 Da), protein N-terminal acetylation (+42.011 Da), and a combination of N-terminal methionine loss and acetylation (+89.030 Da). Static modifications were applied for cysteine carbamidomethylation (+57.021 Da) and TMTpro labeling on the N-terminus or lysines (+304.207 Da). To determine TMT labeling and alkylation efficiencies, the TMTpro and carbamidomethylation modifications were set to dynamic instead of static in Proteome Discover. The proportion of modified vs. unmodified PSMs were then determined.

To control false discoveries, the peptide-level false discovery rate (FDR) was set to 1% using the Percolator algorithm. Quantification of TMT reporter ions was performed by including only those with an average signal-to-noise ratio greater than 10:1 and a co-isolation percentage of less than 25%. TMT reporter ion intensities were summed for peptides assigned to the same protein, including razor peptides, which are shared by multiple proteins but assigned to the one providing the most confident identification. Protein identifications were filtered at a 1% FDR threshold, and protein grouping was conducted according to the parsimony principle, which minimizes the number of protein groups assigned to each peptide.

Median normalization was subsequently carried out using custom R code available on GitHub (https://github.com/Plate-Lab/). Briefly, correction factors per TMT channel were applied to the raw abundances of each observation in the corresponding channel. The mass spectrometry proteomics data are deposited to the ProteomeXchange Consortium via the PRIDE partner repository under the accession code PXD060786. All other necessary data are contained within the manuscript or can be shared by the Lead Contact upon request.

## Abbreviations

(TMT): Tandem Mass Tags
(SP3): Single-Pot Solid-Phase-Enhanced Sample Preparation
(UPR): Unfolded Protein Response
(MS): mass spectrometry
(CV): coefficient of variation
(LC): liquid chromatography
(IAA): iodoacetamide
(TCEP): tris(2-carboxyethyl)phosphine
(TMP): trimethoprim
(DOX): doxycycline
(MudPIT): Multidimensional Protein Identification Technology
(DDA): data dependent acquisition
(PBS): phosphate buffered saline
(EDTA): ethylenediaminetetraacetic acid
(RIPA): radioimmunoprecipitation assay
(ACN): acetonitrile
(FA): formic acid
(FDR): false discovery rate
(FKBP): two FK506-Binding Protein
(dDHFR): destabilized DHFR
(HEPA): High Efficiency Particulate Air.

## Supporting Information

The following files are available free of charge.

- Verification of UPR activation via western blot (Supporting Figure 1); Identification of UPR markers for XBP1s and PERK 18 Hr comparing biomek and manual (Supporting Figure 2) (DOC)
- Estimation of hands-on time vs waiting periods for different sample preparation times (Table S1); Proteome discoverer search file for manual sample preparation (Table S2); Proteome discoverer search file for Biomek sample preparation (Table S3) (xlsx)
- Example of Biomek deck layouts during 96 sample preparation (Supporting Figure 3) (PDF)

## Author Contributions

Conceptualization, L.A.B., J.N.H., L.P.; Investigation, L.A.B, J.N.H.; Writing – Original Draft, L.A.B., J.N.H.; Writing – Review & Editing, L.A.B., J.N.H., L.P.; Visualization, L.A.B., J.N.H.; Supervision, L.P.; Funding Acquisition, L.P.

## Funding Sources

This work was funded by R35GM133552 (National Institute of General Medical Sciences) and Vanderbilt University start-up funds.

## Acknowledgements

None.

## Notes

### Competing Interest Statement

The authors have declared no competing interest.

