## Supporting Information for "Development of an Adaptive, Economical, and Easy-to-Use SP3-TMT Automated Sample Preparation Workflow for Quantitative Proteomics"

**Supplemental Tables.**

1. Comparison of automated (Accelerome/ Biomek i5) and manual protocol completion times.
2. Proteins identified via the manual preparation.
3. Proteins identified via the Biomek i5 preparation.

**Supplemental Figures.**

1. Western blot validation of activated HEK$^{293}$DAX and Fv2e-PERK cells.
2. XBP1s/ PERK target protein quantification, coefficient percentages and correlation plot analysis.

**Supplemental Files.**

1. Deck setups for Biomek sample preparation (pdf).

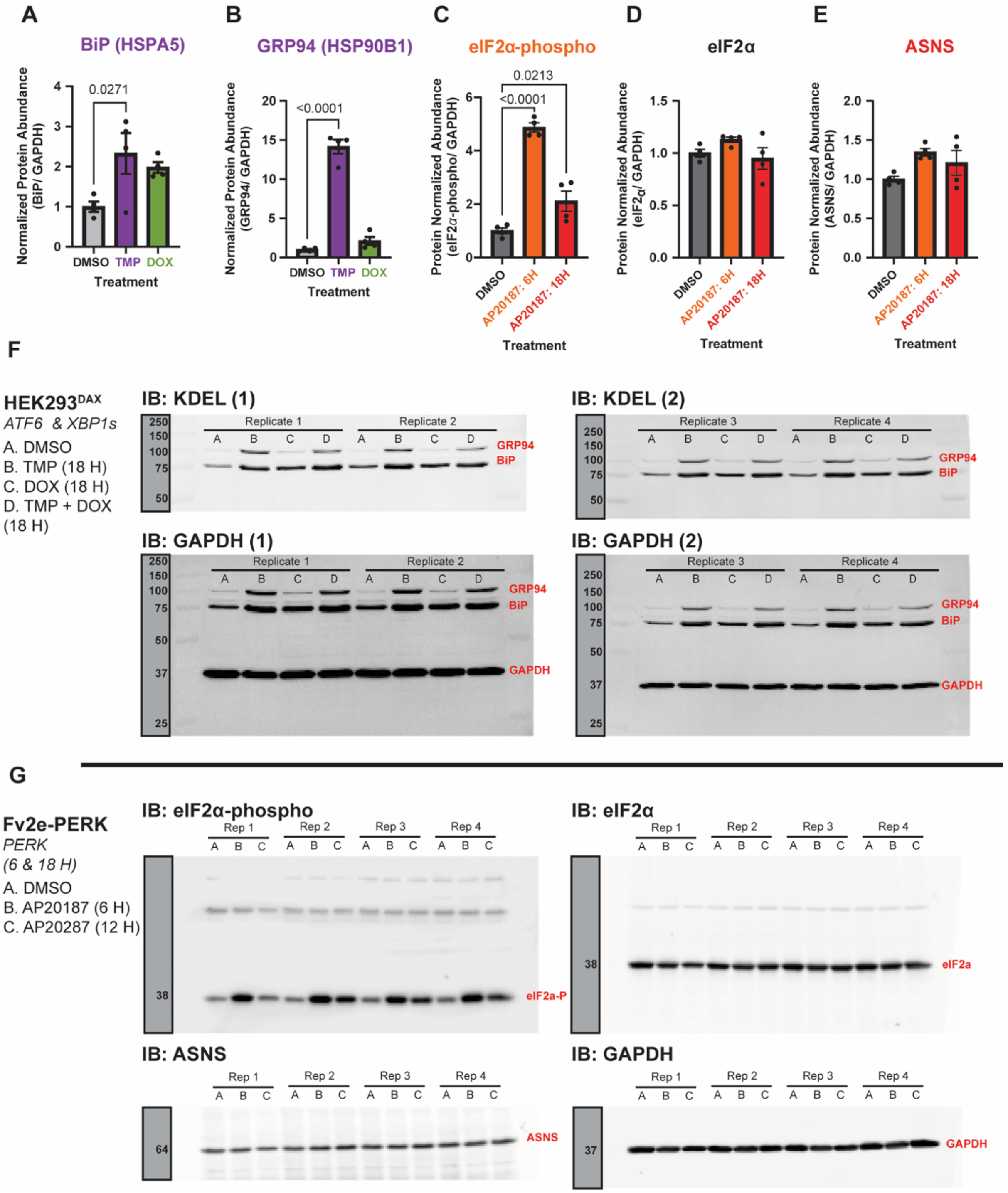


**Supplemental Fig 1. Western blot validation of activated HEK[293]DAX and Fv2e-PERK cell activation.** To upregulate the transcription factors, ATF6 and XBP1s, HEK[293]DAX cells were treated with TMP (10 µM) and DOX (1 µg/mL), respectively for 16 hours. **(A/B)** BiP and HSP90B1 target proteins were utilized to validate activation of ATF6. **(C/D)** Fv2e-PERK cells were alternatively treated with AP20187 (5 nM) for 6 and 18 hours resulting in the phosphorylation of eIF2α (visualized via western blot). **(E)** ASNS was additionally utilized to probe for PERK activation. **(F)** Western blots for HEK[293] DAX cell line activation of ATF6 and XBP1s. **(G)** Western blots of Fv2e-PERK cell activation for 6 and 18 hours.

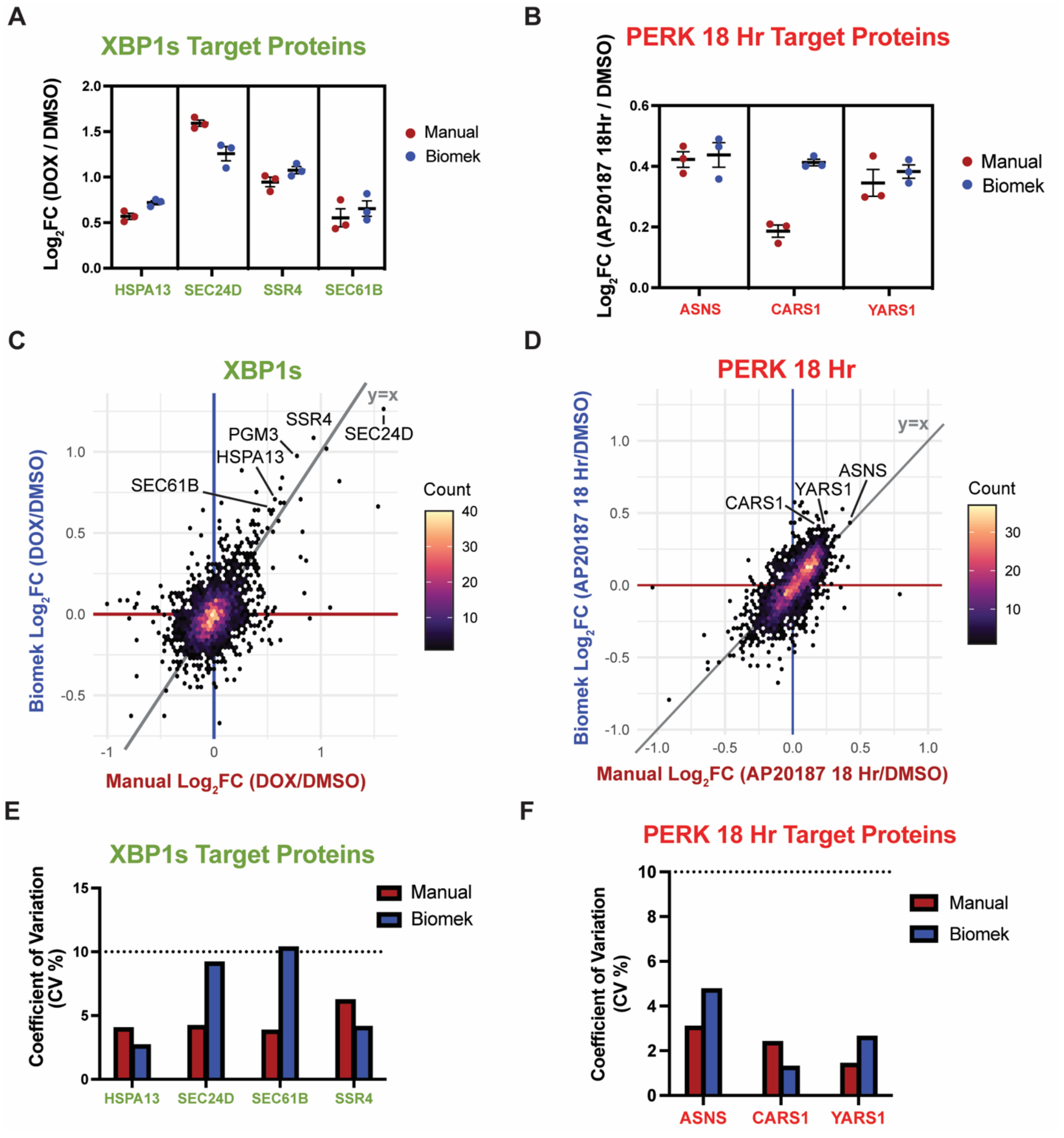


**Supplemental Figure 2. Identification of core UPR markers for both XBP1s and PERK 18 Hr in the Biomek and manual sample preparation.** $Log_2$ fold change of core **(A)** XBP1s and **(B)** PERK 18 Hr between the two methods. Correlation plot of $Log_2$ fold change of **(C)** XBP1s and **(D)** PERK 18 Hr comparing the two sample preparations. Coefficient of variation of select **(E)** XBP1s target and select **(F)** PERK targets.
