## Supporting Figure 3 for "Development of an Adaptive, Economical, and Easy-to-Use SP3-TMT Automated Sample Preparation Workflow for Quantitative Proteomics"

### Labware Setup

This report shows where to place labware on the deck. The quantities and types of labware are listed below.
Start Time: (Print Time: 1/13/2025 7:50:19 PM)

**Method:** GFP Pull Down_V1.1_LowVol2_21Jan2024, **Project:** Biomek i5 MC

#### Overall Setup Information

The left pod should have no tips loaded.

##### Tips and Empty Labware Deck Setup

Ensure that labware is placed on the deck in the positions below.

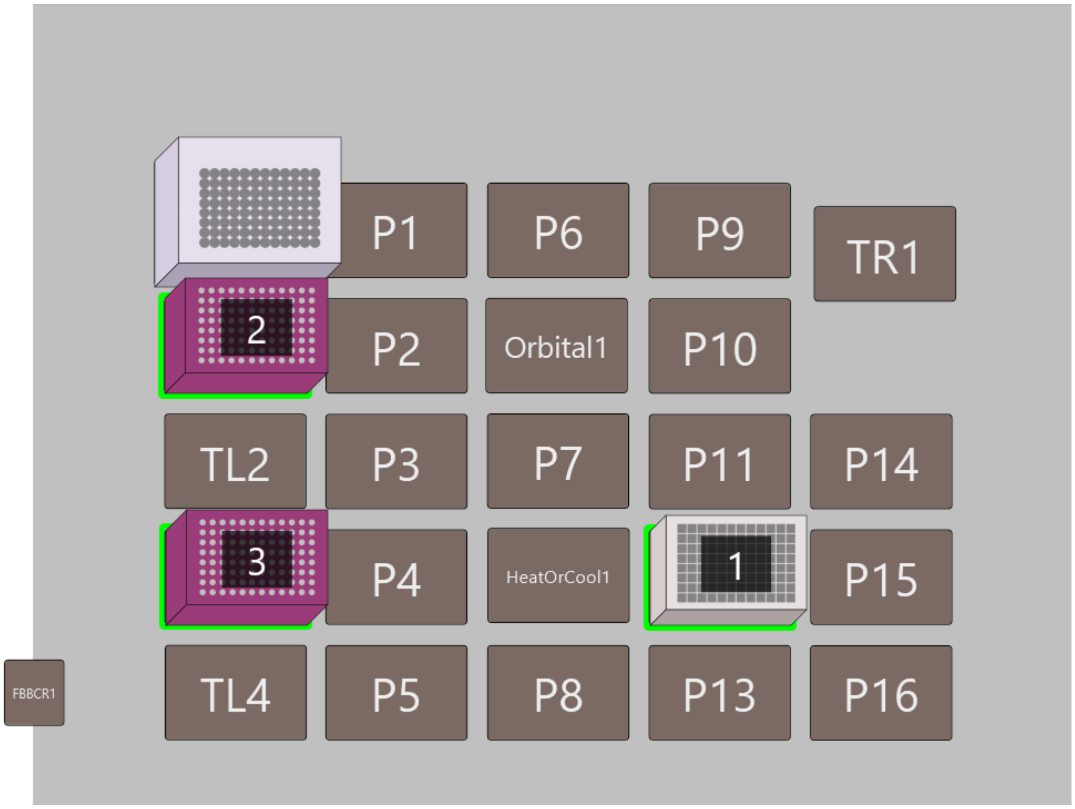


| # | Name | Type | Position | Stack Depth |
|---|---|---|---|---|
| 1 | Dest | ab_0932 | P12 | 1 |
| 2 | Water_Tips | BC90 | TL1 | 1 |
| 3 | Sample_Tips | BC90 | TL3 | 1 |

#### 2. Samples and Reagents

##### Labware Setup for Sample (Bio_RadPCR96)

| | 1 | 2 | 3 | 4 | 5 | 6 | 7 | 8 | 9 | 10 | 11 | 12 |
|---|---|---|---|---|---|---|---|---|---|---|---|---|
| A | Sample 45 uL | Sample 45 uL | Sample 45 uL | Sample 45 uL | Sample 45 uL | Sample 45 uL | Sample 45 uL | Sample 45 uL | Sample 45 uL | Sample 45 uL | Sample 45 uL | Sample 45 uL |
| B | Sample 45 uL | Sample 45 uL | Sample 45 uL | Sample 45 uL | Sample 45 uL | Sample 45 uL | Sample 45 uL | Sample 45 uL | Sample 45 uL | Sample 45 uL | Sample 45 uL | Sample 45 uL |
| C | Sample 45 uL | Sample 45 uL | Sample 45 uL | Sample 45 uL | Sample 45 uL | Sample 45 uL | Sample 45 uL | Sample 45 uL | Sample 45 uL | Sample 45 uL | Sample 45 uL | Sample 45 uL |
| D | Sample 45 uL | Sample 45 uL | Sample 45 uL | Sample 45 uL | Sample 45 uL | Sample 45 uL | Sample 45 uL | Sample 45 uL | Sample 45 uL | Sample 45 uL | Sample 45 uL | Sample 45 uL |
| E | Sample 45 uL | Sample 45 uL | Sample 45 uL | Sample 45 uL | Sample 45 uL | Sample 45 uL | Sample 45 uL | Sample 45 uL | Sample 45 uL | Sample 45 uL | Sample 45 uL | Sample 45 uL |
| F | Sample 45 uL | Sample 45 uL | Sample 45 uL | Sample 45 uL | Sample 45 uL | Sample 45 uL | Sample 45 uL | Sample 45 uL | Sample 45 uL | Sample 45 uL | Sample 45 uL | Sample 45 uL |
| G | Sample 45 uL | Sample 45 uL | Sample 45 uL | Sample 45 uL | Sample 45 uL | Sample 45 uL | Sample 45 uL | Sample 45 uL | Sample 45 uL | Sample 45 uL | Sample 45 uL | Sample 45 uL |
| H | Sample 45 uL | Sample 45 uL | Sample 45 uL | Sample 45 uL | Sample 45 uL | Sample 45 uL | Sample 45 uL | Sample 45 uL | Sample 45 uL | Sample 45 uL | Sample 45 uL | Sample 45 uL |

Place Sample(Bio_RadPCR96) on position P13

#### Samples and Reagents Deck Setup

Ensure that labware is placed on the deck in the positions below.

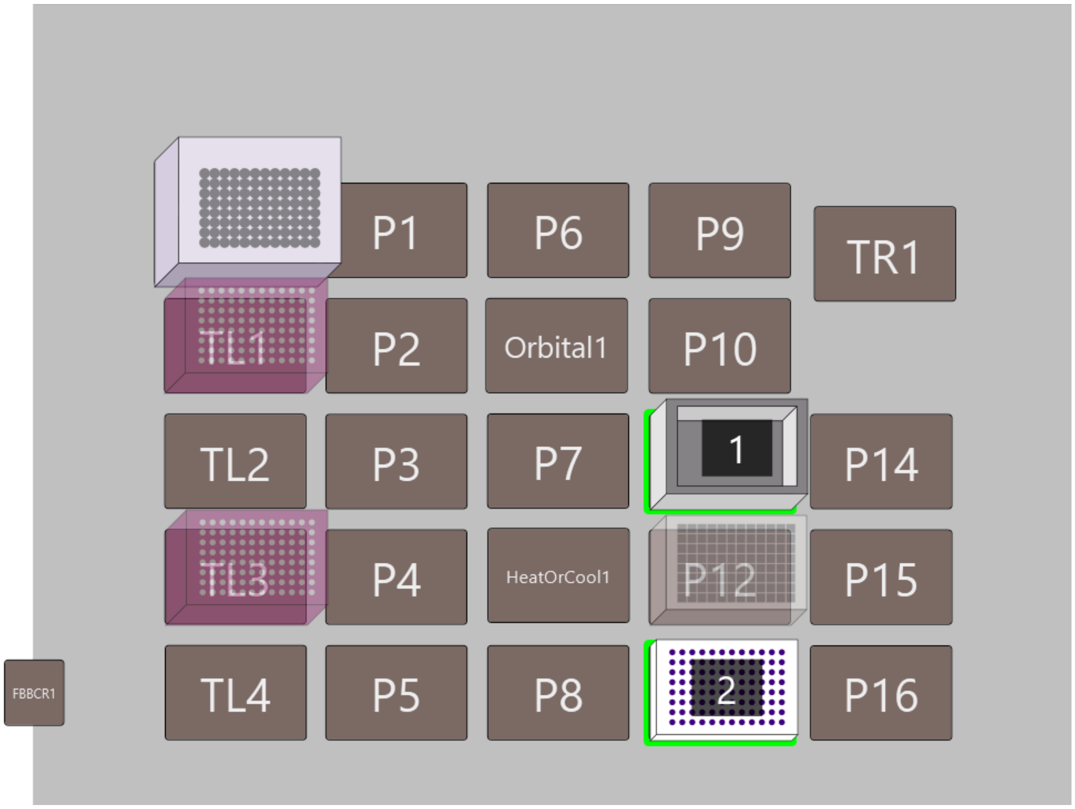

| # | Name | Type | Position | Stack Depth |
|---|---|---|---|---|
| 1 | Water | AgilentReservoir | P11 | 1 |
| 2 | Sample | Bio_RadPCR96 | P13 | 1 |

### Final Deck Setup

Below is the final deck layout. Ensure that all the highlighted labware are in the specified positions.

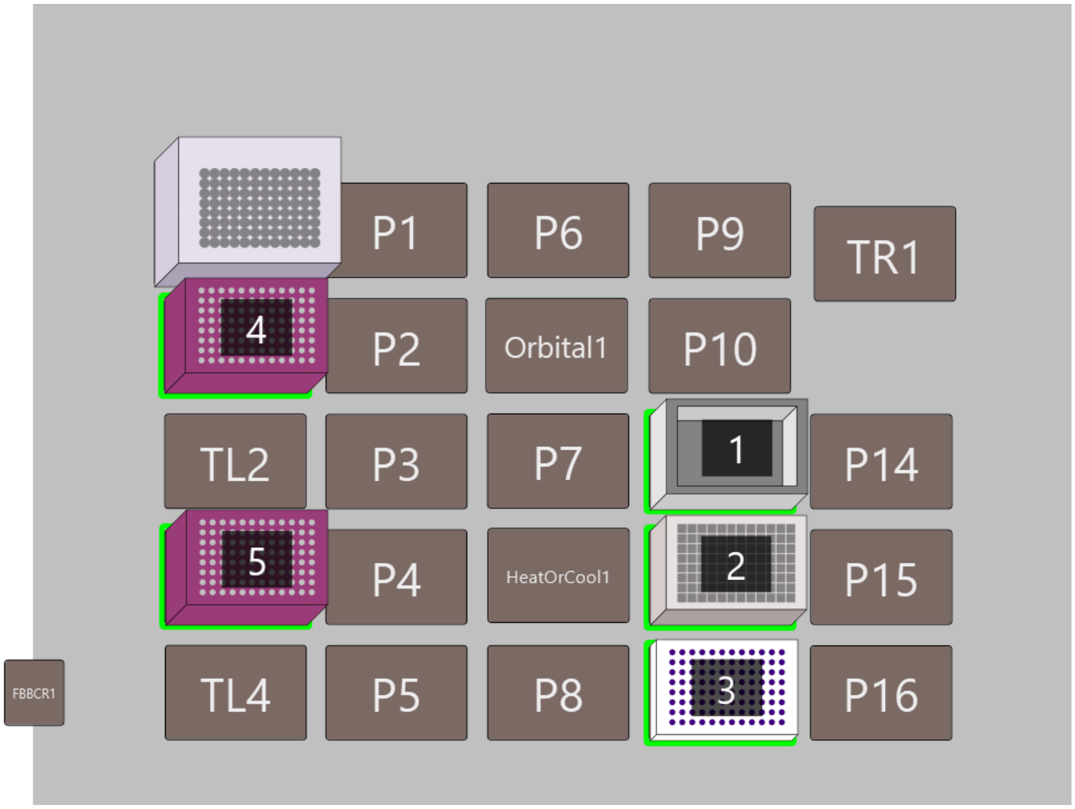


| # | Name | Type | Position | Stack Depth |
|---|---|---|---|---|
| 1 | Water | AgilentReservoir | P11 | 1 |
| 2 | Dest | ab_0932 | P12 | 1 |
| 3 | Sample | Bio_RadPCR96 | P13 | 1 |
| 4 | Water_Tips | BC90 | TL1 | 1 |
| 5 | Sample_Tips | BC90 | TL3 | 1 |

### Method Details

### Labware Setup

This report shows where to place labware on the deck. The quantities and types of labware are listed below.
Start Time:  (Print Time: 1/14/2025 10:21:40 AM)

**Method:** SP3 Bead_V1.1_Pauses_50ul, **Project:** Biomek i5 MC

### Overall Setup Information

The left pod should have no tips loaded.

### 1. New Labware Group

#### Labware Setup for Sample (ab_0932)

| | 1 | 2 | 3 | 4 | 5 | 6 | 7 | 8 | 9 | 10 | 11 | 12 |
|---|---|---|---|---|---|---|---|---|---|---|---|---|
| A | 20 ug protein 70 uL | 20 ug protein 70 uL | 20 ug protein 70 uL | 20 ug protein 70 uL | 20 ug protein 70 uL | 20 ug protein 70 uL | 20 ug protein 70 uL | 20 ug protein 70 uL | 20 ug protein 70 uL | 20 ug protein 70 uL | 20 ug protein 70 uL | 20 ug protein 70 uL |
| B | 20 ug protein 70 uL | 20 ug protein 70 uL | 20 ug protein 70 uL | 20 ug protein 70 uL | 20 ug protein 70 uL | 20 ug protein 70 uL | 20 ug protein 70 uL | 20 ug protein 70 uL | 20 ug protein 70 uL | 20 ug protein 70 uL | 20 ug protein 70 uL | 20 ug protein 70 uL |
| C | 20 ug protein 70 uL | 20 ug protein 70 uL | 20 ug protein 70 uL | 20 ug protein 70 uL | 20 ug protein 70 uL | 20 ug protein 70 uL | 20 ug protein 70 uL | 20 ug protein 70 uL | 20 ug protein 70 uL | 20 ug protein 70 uL | 20 ug protein 70 uL | 20 ug protein 70 uL |
| D | 20 ug protein 70 uL | 20 ug protein 70 uL | 20 ug protein 70 uL | 20 ug protein 70 uL | 20 ug protein 70 uL | 20 ug protein 70 uL | 20 ug protein 70 uL | 20 ug protein 70 uL | 20 ug protein 70 uL | 20 ug protein 70 uL | 20 ug protein 70 uL | 20 ug protein 70 uL |
| E | 20 ug protein 70 uL | 20 ug protein 70 uL | 20 ug protein 70 uL | 20 ug protein 70 uL | 20 ug protein 70 uL | 20 ug protein 70 uL | 20 ug protein 70 uL | 20 ug protein 70 uL | 20 ug protein 70 uL | 20 ug protein 70 uL | 20 ug protein 70 uL | 20 ug protein 70 uL |
| F | 20 ug protein 70 uL | 20 ug protein 70 uL | 20 ug protein 70 uL | 20 ug protein 70 uL | 20 ug protein 70 uL | 20 ug protein 70 uL | 20 ug protein 70 uL | 20 ug protein 70 uL | 20 ug protein 70 uL | 20 ug protein 70 uL | 20 ug protein 70 uL | 20 ug protein 70 uL |
| G | 20 ug protein 70 uL | 20 ug protein 70 uL | 20 ug protein 70 uL | 20 ug protein 70 uL | 20 ug protein 70 uL | 20 ug protein 70 uL | 20 ug protein 70 uL | 20 ug protein 70 uL | 20 ug protein 70 uL | 20 ug protein 70 uL | 20 ug protein 70 uL | 20 ug protein 70 uL |
| H | 20 ug protein 70 uL | 20 ug protein 70 uL | 20 ug protein 70 uL | 20 ug protein 70 uL | 20 ug protein 70 uL | 20 ug protein 70 uL | 20 ug protein 70 uL | 20 ug protein 70 uL | 20 ug protein 70 uL | 20 ug protein 70 uL | 20 ug protein 70 uL | 20 ug protein 70 uL |

Place Sample(ab_0932) on position P11

#### Labware Setup for Reagents (ab_0932)

| | 1 | 2 | 3 | 4 | 5 | 6 | 7 | 8 | 9 | 10 | 11 | 12 |
|---|---|---|---|---|---|---|---|---|---|---|---|---|
| A | 100 mM DTT 113.8uL | 200 mM IAA 133uL | | | | | | | | | | |
| B | 100 mM DTT 113.8uL | 200 mM IAA 133uL | | | | | | | | | | |
| C | 100 mM DTT 113.8uL | 200 mM IAA 133uL | | | | | | | | | | |
| D | 100 mM DTT 113.8uL | 200 mM IAA 133uL | | | | | | | | | | |
| E | 100 mM DTT 113.8uL | 200 mM IAA 133uL | | | | | | | | | | |
| F | 100 mM DTT 113.8uL | 200 mM IAA 133uL | | | | | | | | | | |
| G | 100 mM DTT 113.8uL | 200 mM IAA 133uL | | | | | | | | | | |
| H | 100 mM DTT 113.8uL | 200 mM IAA 133uL | | | | | | | | | | |

Place Reagents(ab_0932) on position P12

#### New Labware Group Deck Setup

Ensure that labware is placed on the deck in the positions below.

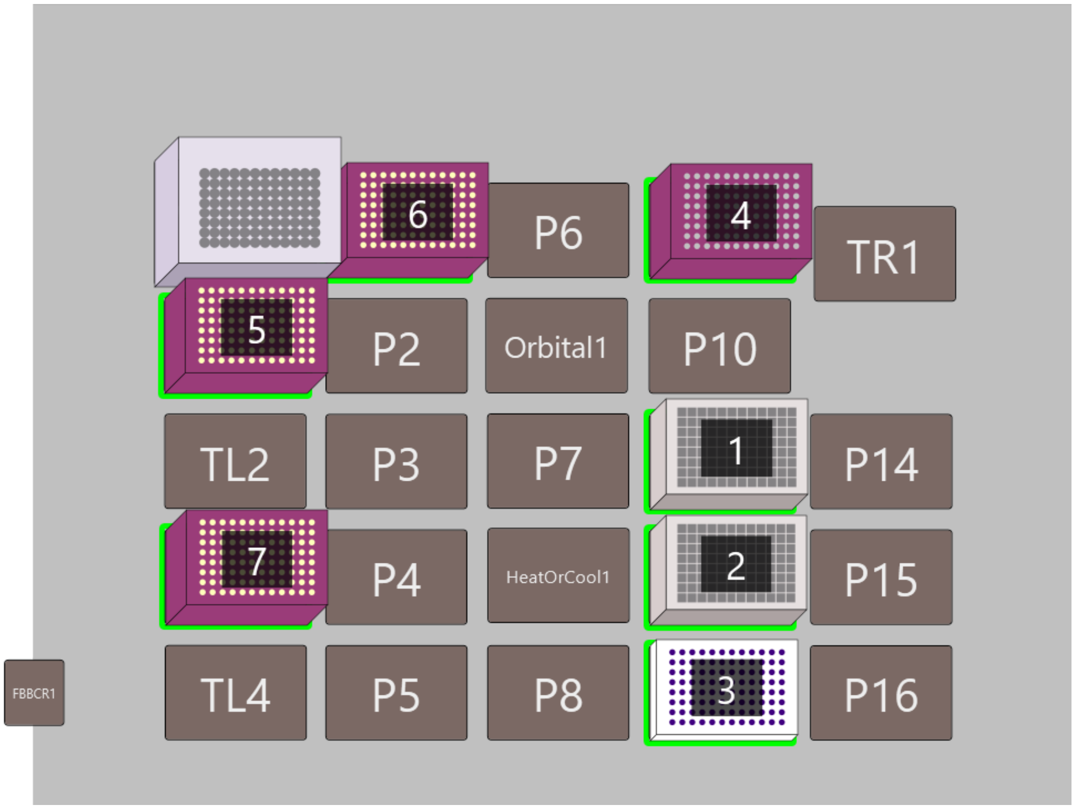

| # | Name | Type | Position | Stack Depth |
|---|---|---|---|---|
| 1 | Sample | ab_0932 | P11 | 1 |
| 2 | Reagents | ab_0932 | P12 | 1 |
| 3 | Gel | Bio_RadPCR96 | P13 | 1 |
| 4 | Gel_Tips | BC90 | P9 | 1 |
| 5 | DTT_Tips1 | BC50F | TL1 | 1 |
| 6 | DTT_Tips2 | BC50F | P1 | 1 |
| 7 | IAA_Tips | BC50F | TL3 | 1 |

### Final Deck Setup

Below is the final deck layout. Ensure that all the highlighted labware are in the specified positions.

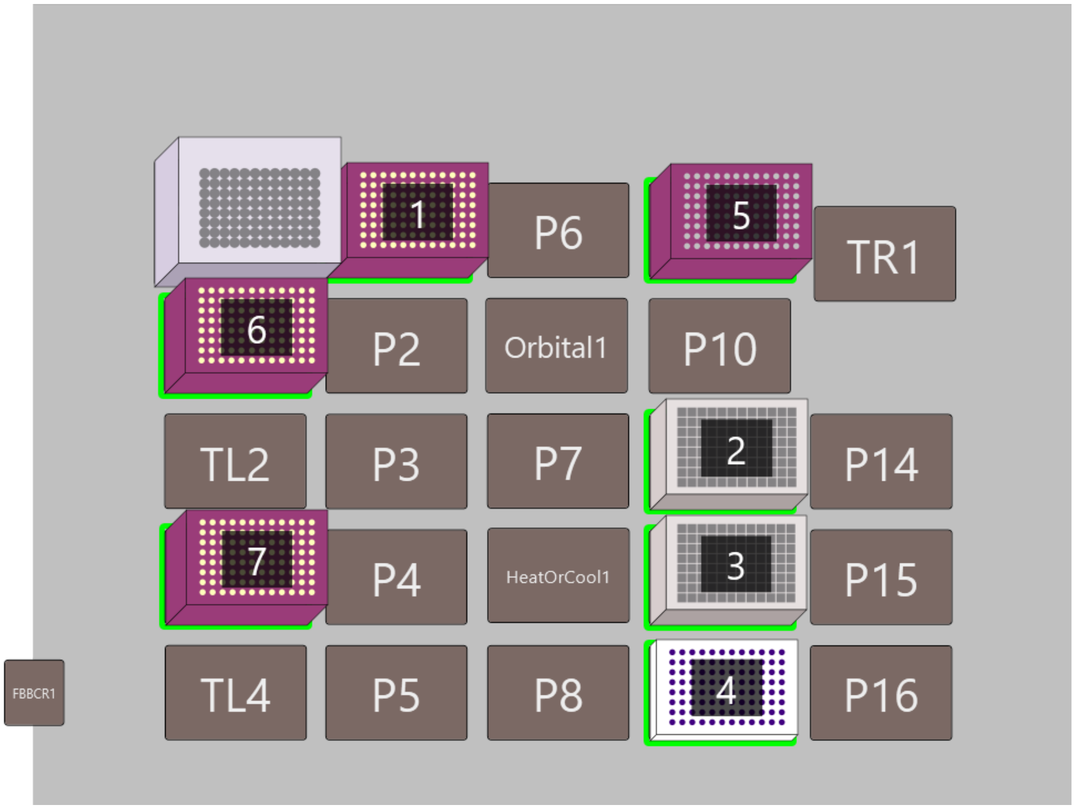

### Labware Setup

This report shows where to place labware on the deck. The quantities and types of labware are listed below.
Start Time: (Print Time: 1/14/2025 10:52:50 AM)

**Method:** SP3 Bead_V1.1_Pauses_50ul, **Project:** Biomek i5 MC

#### Overall Setup Information

The left pod should have no tips loaded.

#### 1. New Labware Group

##### Labware Setup for EtOH (AgilentReservoir)

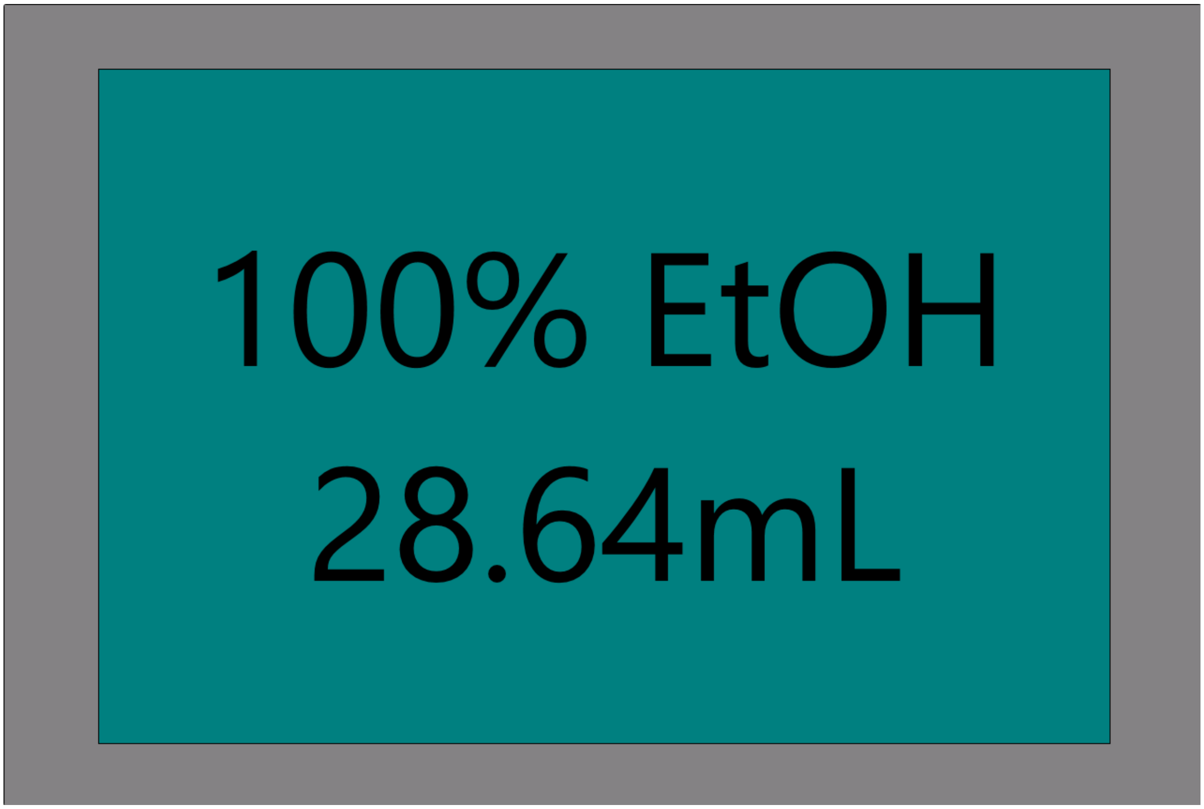


Place EtOH(AgilentReservoir) on position P8

##### Labware Setup for 80% EtOH (AgilentReservoir)

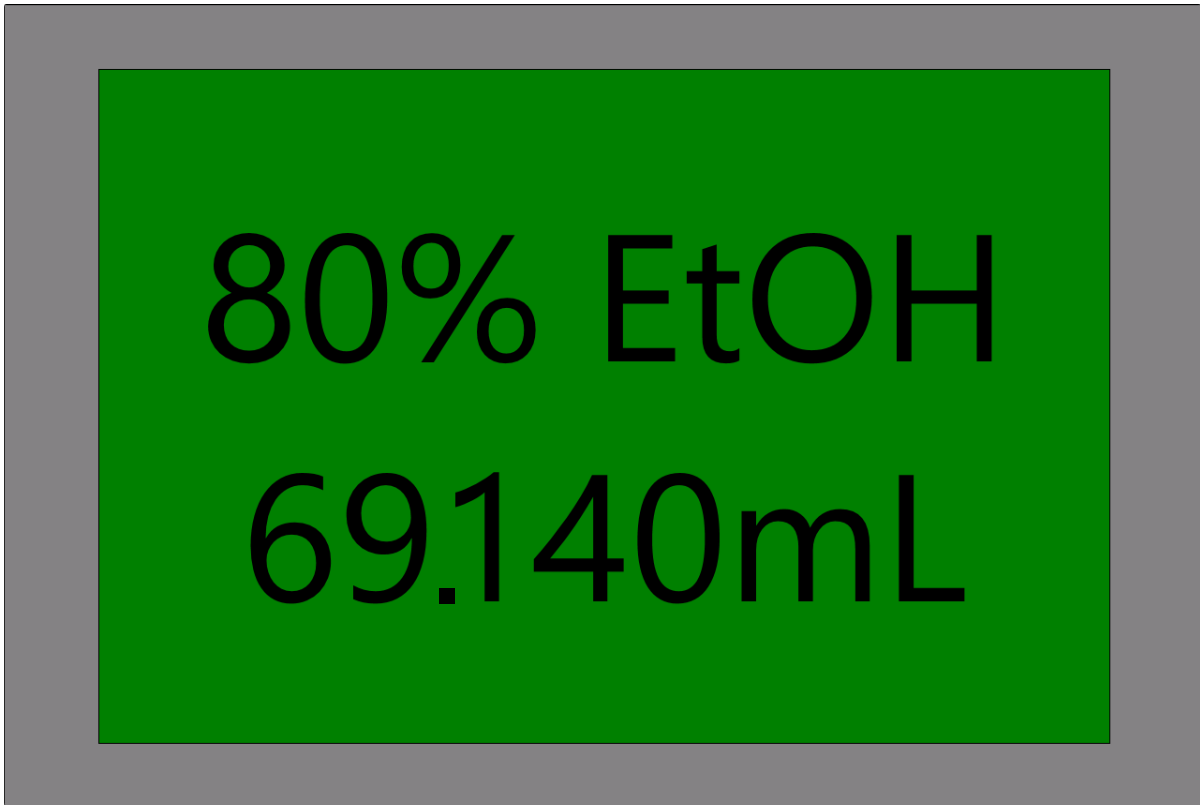


Place 80% EtOH(AgilentReservoir) on position P3

#### Labware Setup for Trypsin (ab_0932)

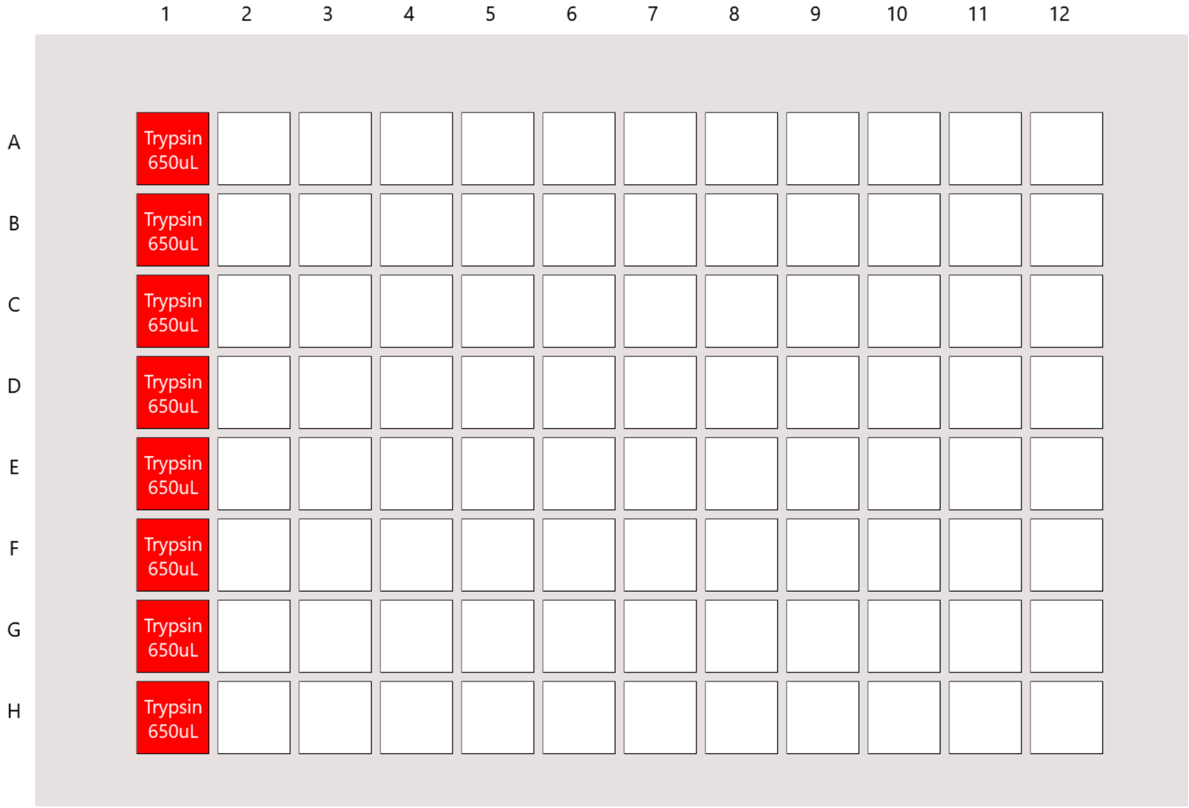


Place Trypsin(ab_0932) on position P10

### Labware Setup for Beads (ab_0932)

Notes:
Only need to put beads where samples are in the sample plate

| | 1 | 2 | 3 | 4 | 5 | 6 | 7 | 8 | 9 | 10 | 11 | 12 |
|---|---|---|---|---|---|---|---|---|---|---|---|---|
| A | Beads 8 uL | Beads 8 uL | Beads 8 uL | Beads 8 uL | Beads 8 uL | Beads 8 uL | Beads 8 uL | Beads 8 uL | Beads 8 uL | Beads 8 uL | Beads 8 uL | Beads 8 uL |
| B | Beads 8 uL | Beads 8 uL | Beads 8 uL | Beads 8 uL | Beads 8 uL | Beads 8 uL | Beads 8 uL | Beads 8 uL | Beads 8 uL | Beads 8 uL | Beads 8 uL | Beads 8 uL |
| C | Beads 8 uL | Beads 8 uL | Beads 8 uL | Beads 8 uL | Beads 8 uL | Beads 8 uL | Beads 8 uL | Beads 8 uL | Beads 8 uL | Beads 8 uL | Beads 8 uL | Beads 8 uL |
| D | Beads 8 uL | Beads 8 uL | Beads 8 uL | Beads 8 uL | Beads 8 uL | Beads 8 uL | Beads 8 uL | Beads 8 uL | Beads 8 uL | Beads 8 uL | Beads 8 uL | Beads 8 uL |
| E | Beads 8 uL | Beads 8 uL | Beads 8 uL | Beads 8 uL | Beads 8 uL | Beads 8 uL | Beads 8 uL | Beads 8 uL | Beads 8 uL | Beads 8 uL | Beads 8 uL | Beads 8 uL |
| F | Beads 8 uL | Beads 8 uL | Beads 8 uL | Beads 8 uL | Beads 8 uL | Beads 8 uL | Beads 8 uL | Beads 8 uL | Beads 8 uL | Beads 8 uL | Beads 8 uL | Beads 8 uL |
| G | Beads 8 uL | Beads 8 uL | Beads 8 uL | Beads 8 uL | Beads 8 uL | Beads 8 uL | Beads 8 uL | Beads 8 uL | Beads 8 uL | Beads 8 uL | Beads 8 uL | Beads 8 uL |
| H | Beads 8 uL | Beads 8 uL | Beads 8 uL | Beads 8 uL | Beads 8 uL | Beads 8 uL | Beads 8 uL | Beads 8 uL | Beads 8 uL | Beads 8 uL | Beads 8 uL | Beads 8 uL |

Place Beads(ab_0932) on position P7

### New Labware Group Deck Setup

Ensure that labware is placed on the deck in the positions below.

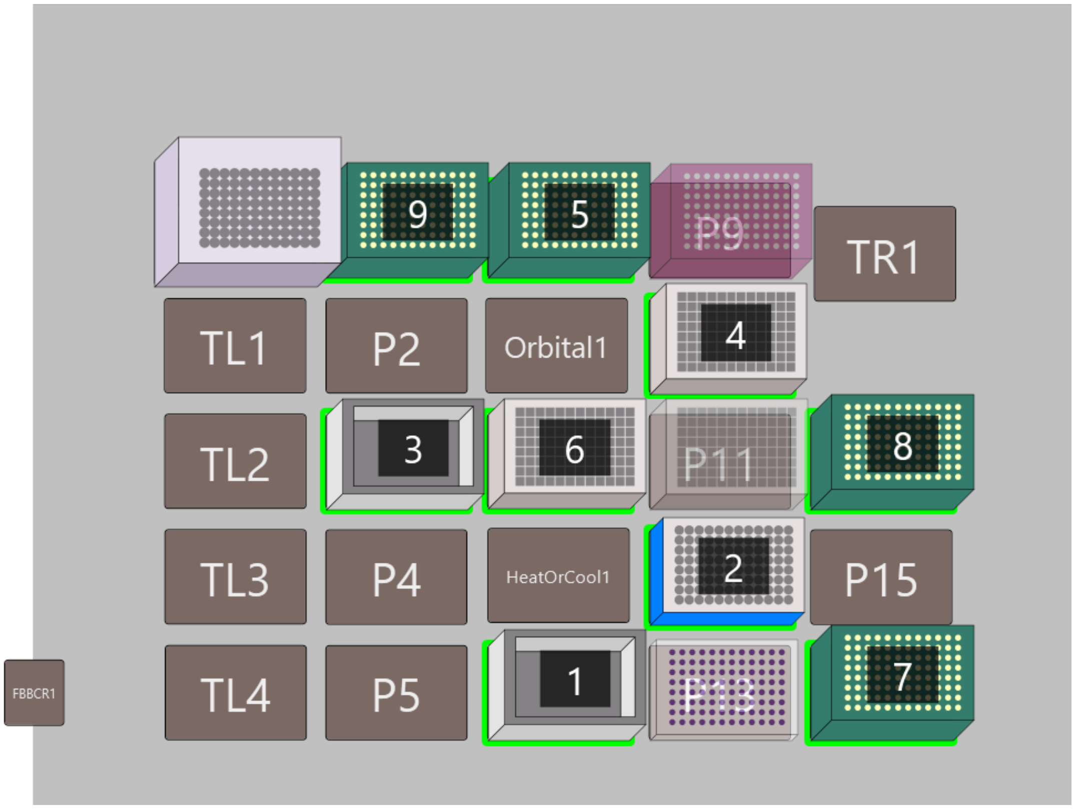


| # | Name | Type | Position | Stack Depth |
|---|---|---|---|---|
| 1 | EtOH | AgilentReservoir | P8 | 1 |
| 2 | Magnet | magnum_flx | P12 | 1 |
| 3 | 80% EtOH | AgilentReservoir | P3 | 1 |
| 4 | Trypsin | ab_0932 | P10 | 1 |
| 5 | Trypsin_Tips | BC190F | P6 | 1 |
| 6 | Beads | ab_0932 | P7 | 1 |
| 7 | Sample_Tips | BC190F | P16 | 1 |
| 8 | EtOH_Tips | BC190F | P14 | 1 |
| 9 | EtOH_Wash | BC190F | P1 | 1 |

### Final Deck Setup

Below is the final deck layout. Ensure that all the highlighted labware are in the specified positions.

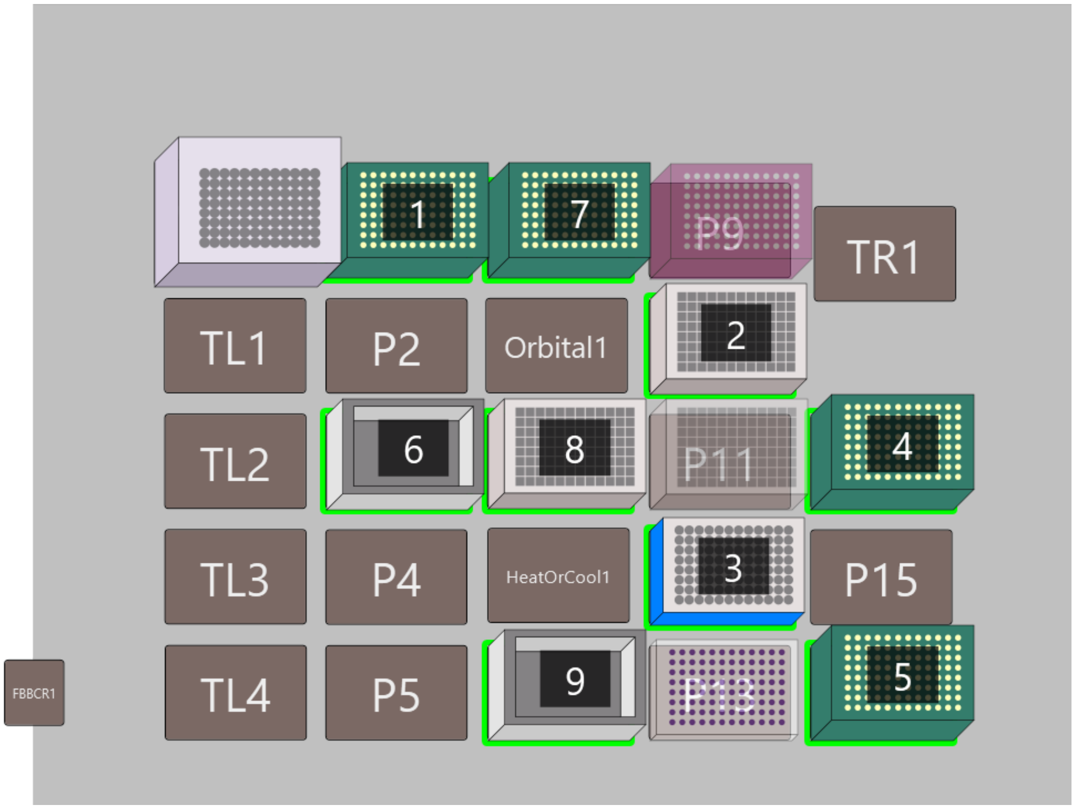


| # | Name | Type | Position | Stack Depth |
|---|---|---|---|---|
| 1 | EtOH_Wash | BC190F | P1 | 1 |
| 2 | Trypsin | ab_0932 | P10 | 1 |
| 3 | Magnet | magnum_flx | P12 | 1 |
| 4 | EtOH_Tips | BC190F | P14 | 1 |
| 5 | Sample_Tips | BC190F | P16 | 1 |
| 6 | 80% EtOH | AgilentReservoir | P3 | 1 |
| 7 | Trypsin_Tips | BC190F | P6 | 1 |
| 8 | Beads | ab_0932 | P7 | 1 |
| 9 | EtOH | AgilentReservoir | P8 | 1 |

### Method Details

### Labware Setup

This report shows where to place labware on the deck. The quantities and types of labware are listed below.
Start Time: (Print Time: 1/14/2025 1:16:17 PM)

**Method:** SP3 Bead_V1.1_Pauses_50ul, **Project:** Biomek i5 MC

#### Overall Setup Information

The left pod should have no tips loaded.

##### New Labware Group Deck Setup

Ensure that labware is placed on the deck in the positions below.

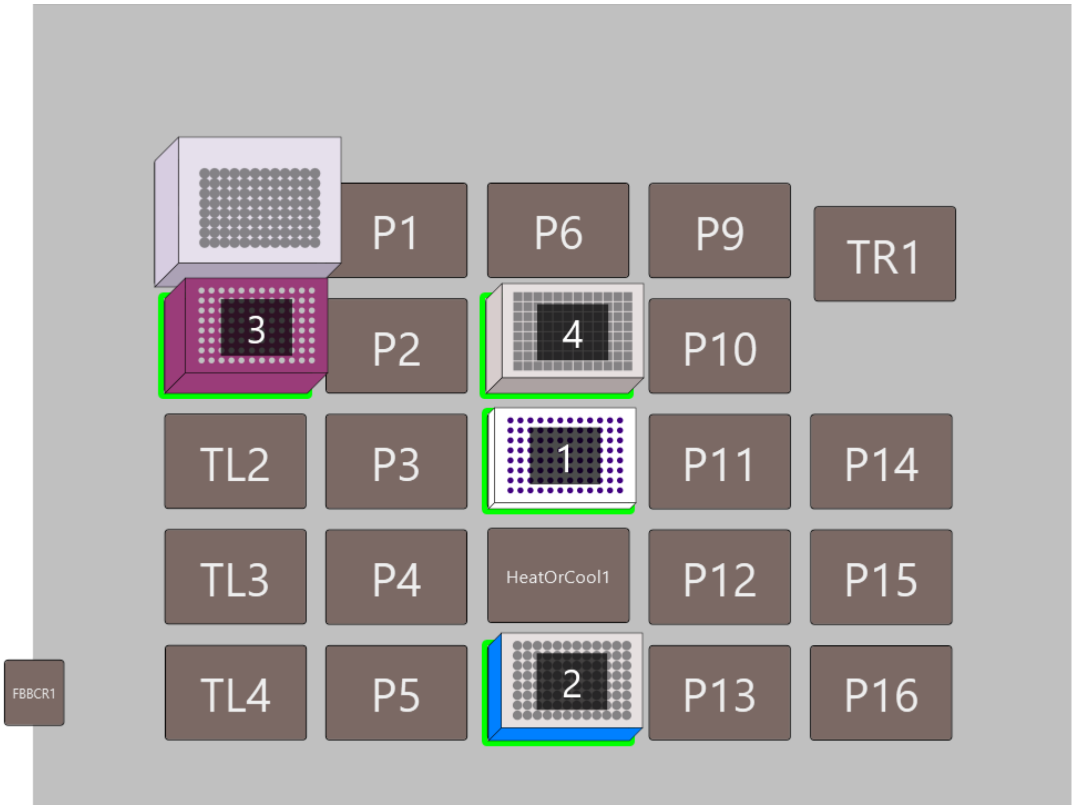


| # | Name | Type | Position | Stack Depth |
|---|---|---|---|---|
| 1 | Supernatent | Bio_RadPCR96 | P7 | 1 |
| 2 | Magnet | magnum_flx | P8 | 1 |
| 3 | Coll_Tips | BC90 | TL1 | 1 |
| 4 | Beads | ab_0932 | Orbital1 | 1 |

#### Final Deck Setup

Below is the final deck layout. Ensure that all the highlighted labware are in the specified positions.

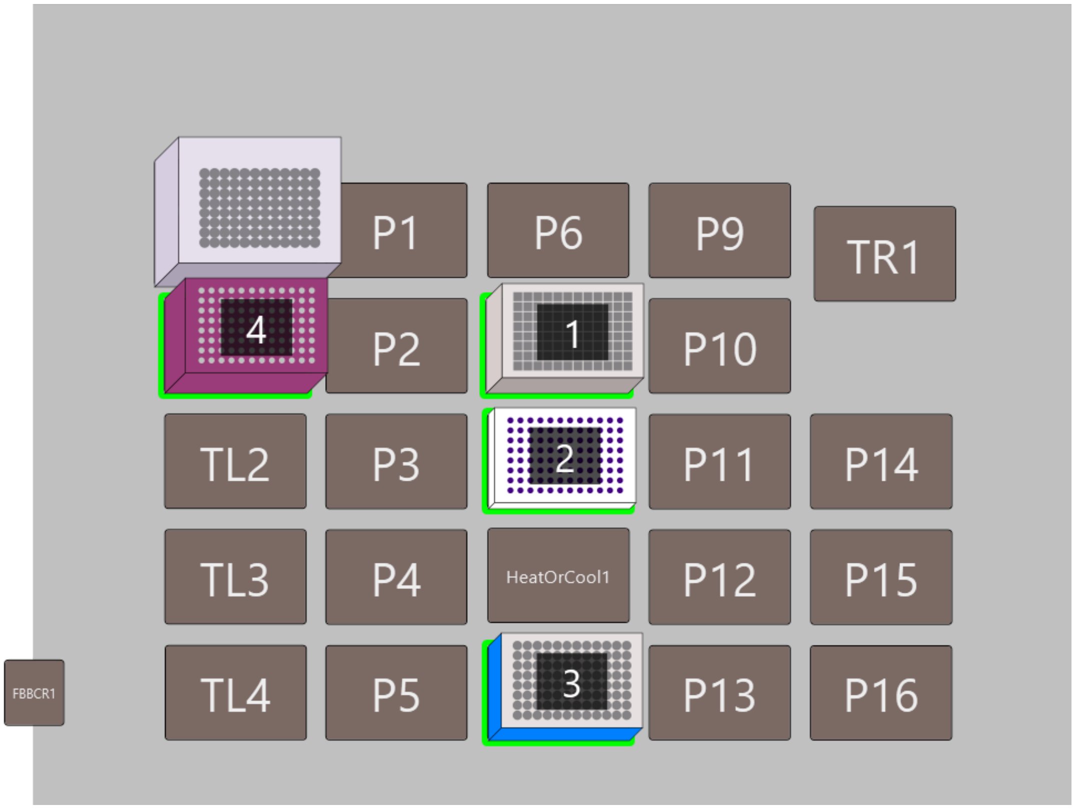


| # | Name | Type | Position | Stack Depth |
|---|---|---|---|---|
| 1 | Beads | ab_0932 | Orbital1 | 1 |
| 2 | Supernatent | Bio_RadPCR96 | P7 | 1 |
| 3 | Magnet | magnum_flx | P8 | 1 |
| 4 | Coll_Tips | BC90 | TL1 | 1 |

### Method Details

### Labware Setup

This report shows where to place labware on the deck. The quantities and types of labware are listed below.
Start Time:  (Print Time: 1/14/2025 1:24:44 PM)

**Method:** TMT Labeling_V1.1, **Project:** Biomek i5 MC

#### Overall Setup Information

The left pod should have no tips loaded.

#### 1. Tips and Empty Labware

##### Labware Setup for Final Tubes (BCTubeBlock_2mlTubes)

Notes:
Add 1 tube for each experiment you are running.

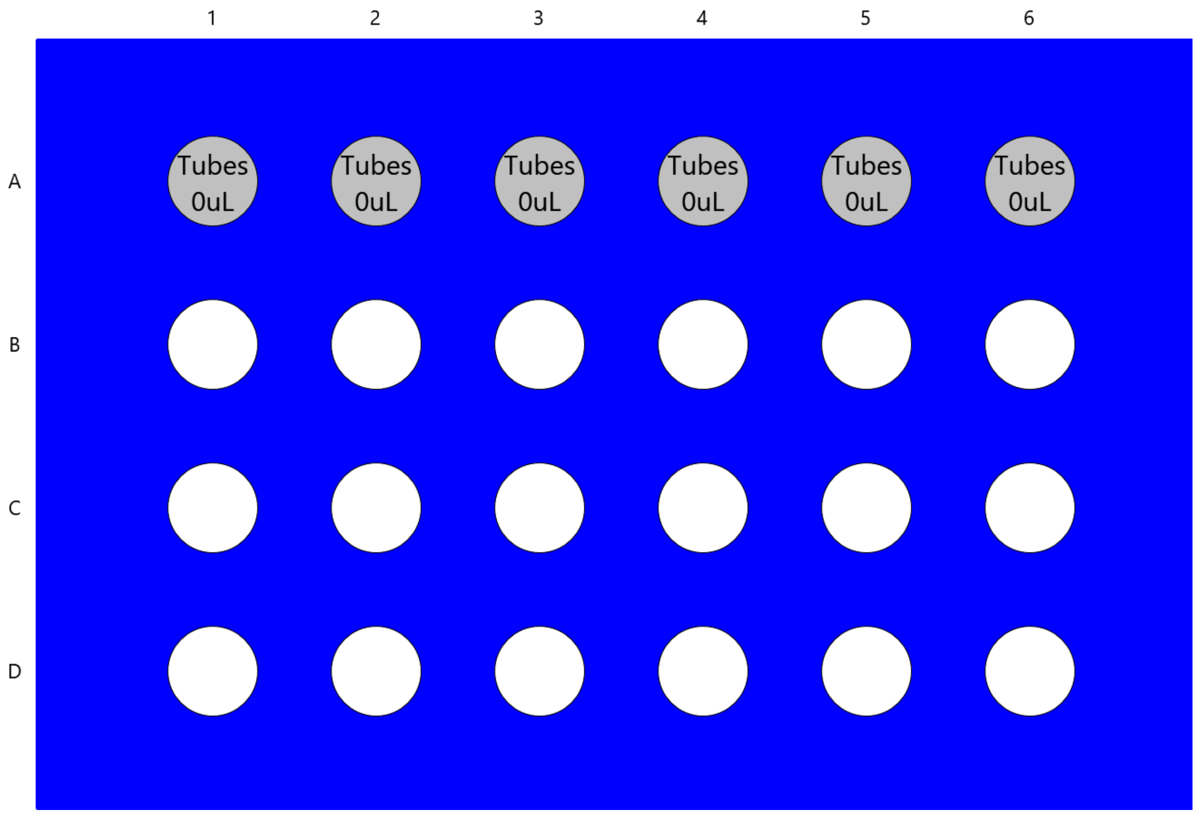


Place Final Tubes(BCTubeBlock_2mlTubes) on position P1

##### Tips and Empty Labware Deck Setup

Ensure that labware is placed on the deck in the positions below.

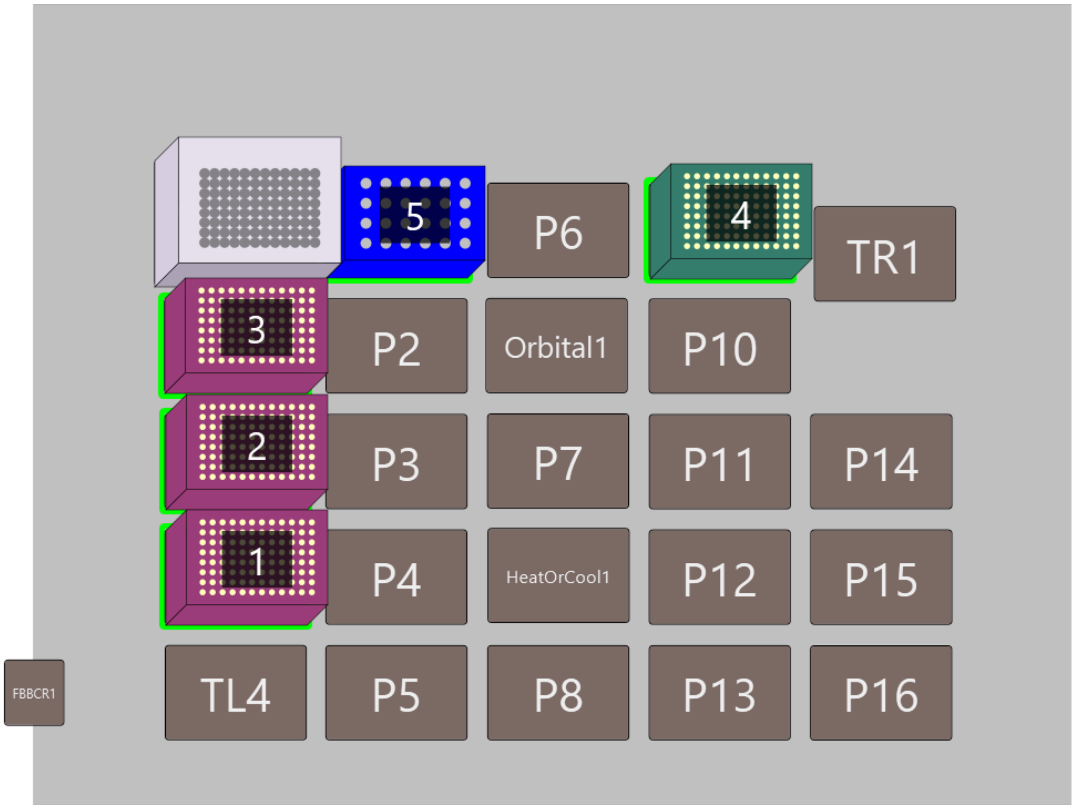


| # | Name | Type | Position | Stack Depth |
|---|---|---|---|---|
| 1 | AmBi Tips | BC50F | TL3 | 1 |
| 2 | TMT Tips | BC50F | TL2 | 1 |
| 3 | Water Tips | BC50F | TL1 | 1 |
| 4 | Pooling Tips | BC190F | P9 | 1 |
| 5 | Final Tubes | BCTubeBlock_2mlTubes | P1 | 1 |

### 2. Reagents

#### Labware Setup for Reagents (ab_0932)

Notes:
Master Mix Composition for 1 reactions:
10uL Hepes + 32.5uL H20uL + .5uL TCEP

| | 1 | 2 | 3 | 4 | 5 | 6 | 7 | 8 | 9 | 10 | 11 | 12 |
|---|---|---|---|---|---|---|---|---|---|---|---|---|
| A | | | Water 145uL | AmBi 73uL | FAcid 611.6uL | | | | | | | |
| B | | | Water 145uL | AmBi 73uL | | | | | | | | |
| C | | | Water 145uL | AmBi 73uL | | | | | | | | |
| D | | | Water 145uL | AmBi 73uL | | | | | | | | |
| E | | | Water 145uL | AmBi 73uL | | | | | | | | |
| F | | | Water 145uL | AmBi 73uL | | | | | | | | |
| G | | | Water 145uL | AmBi 73uL | | | | | | | | |
| H | | | Water 145uL | AmBi 73uL | | | | | | | | |

Place Reagents(ab_0932) on position P8

#### Labware Setup for LVLTubes (LVL_MX500_TMT_Tubes)

| | 1 | 2 | 3 | 4 | 5 | 6 | 7 | 8 | 9 | 10 | 11 | 12 |
|---|---|---|---|---|---|---|---|---|---|---|---|---|
| A | Trypsin 121uL | S1 265uL | S9 265uL | | | | | | | | | |
| B | | S2 265uL | S10 265uL | | | | | | | | | |
| C | | S3 265uL | S11 265uL | | | | | | | | | |
| D | | S4 265uL | S12 265uL | | | | | | | | | |
| E | | S5 265uL | S13 265uL | | | | | | | | | |
| F | | S6 265uL | S14 265uL | | | | | | | | | |
| G | | S7 265uL | S15 265uL | | | | | | | | | |
| H | | S8 265uL | S16 265uL | | | | | | | | | |

Place LVLTubes(LVL_MX500_TMT_Tubes) on position P4

### Labware Setup for Sample (Bio_RadPCR96)

Notes:
Arrange each Experiment in the layout shown Below.

Up to 6 Experiments can fit on one 96 Well Plate

| | 1 | 2 | 3 | 4 | 5 | 6 | 7 | 8 | 9 | 10 | 11 | 12 |
|---|---|---|---|---|---|---|---|---|---|---|---|---|
| A | Sample 50uL | Sample 50uL | Sample 50uL | Sample 50uL | Sample 50uL | Sample 50uL | Sample 50uL | Sample 50uL | Sample 50uL | Sample 50uL | Sample 50uL | Sample 50uL |
| B | Sample 50uL | Sample 50uL | Sample 50uL | Sample 50uL | Sample 50uL | Sample 50uL | Sample 50uL | Sample 50uL | Sample 50uL | Sample 50uL | Sample 50uL | Sample 50uL |
| C | Sample 50uL | Sample 50uL | Sample 50uL | Sample 50uL | Sample 50uL | Sample 50uL | Sample 50uL | Sample 50uL | Sample 50uL | Sample 50uL | Sample 50uL | Sample 50uL |
| D | Sample 50uL | Sample 50uL | Sample 50uL | Sample 50uL | Sample 50uL | Sample 50uL | Sample 50uL | Sample 50uL | Sample 50uL | Sample 50uL | Sample 50uL | Sample 50uL |
| E | Sample 50uL | Sample 50uL | Sample 50uL | Sample 50uL | Sample 50uL | Sample 50uL | Sample 50uL | Sample 50uL | Sample 50uL | Sample 50uL | Sample 50uL | Sample 50uL |
| F | Sample 50uL | Sample 50uL | Sample 50uL | Sample 50uL | Sample 50uL | Sample 50uL | Sample 50uL | Sample 50uL | Sample 50uL | Sample 50uL | Sample 50uL | Sample 50uL |
| G | Sample 50uL | Sample 50uL | Sample 50uL | Sample 50uL | Sample 50uL | Sample 50uL | Sample 50uL | Sample 50uL | Sample 50uL | Sample 50uL | Sample 50uL | Sample 50uL |
| H | Sample 50uL | Sample 50uL | Sample 50uL | Sample 50uL | Sample 50uL | Sample 50uL | Sample 50uL | Sample 50uL | Sample 50uL | Sample 50uL | Sample 50uL | Sample 50uL |

Place Sample(Bio_RadPCR96) on position P7

#### Reagents Deck Setup

Ensure that labware is placed on the deck in the positions below.

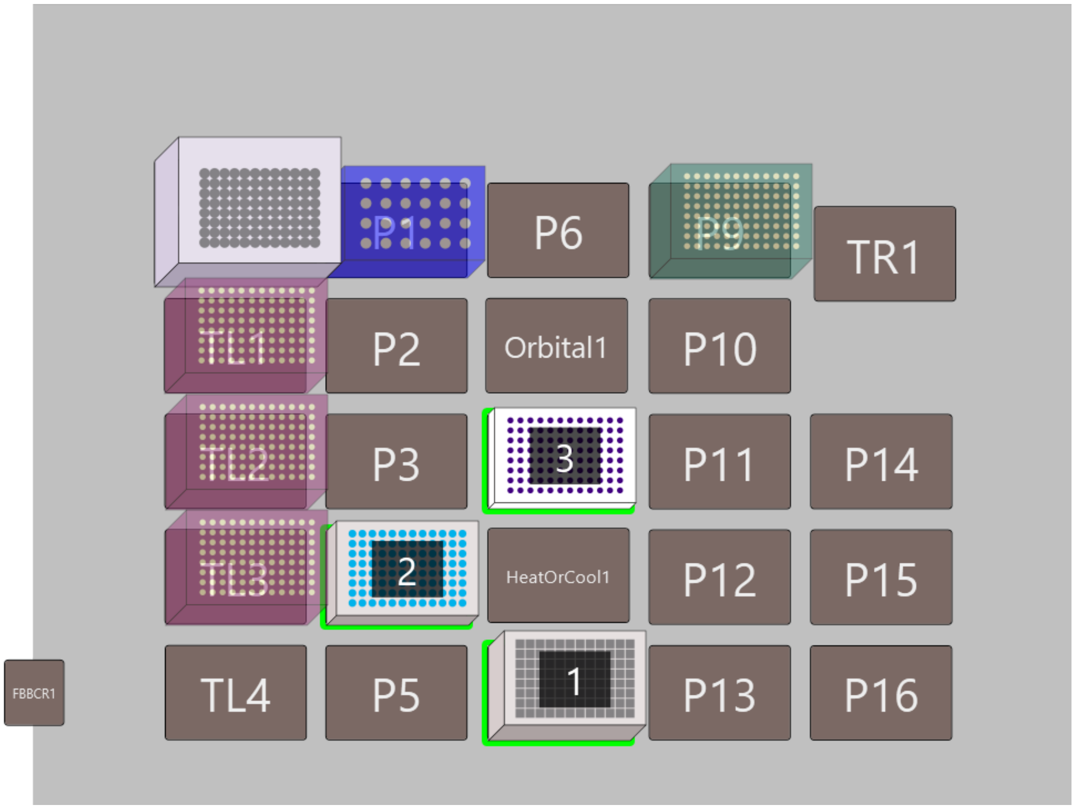


| # | Name | Type | Position | Stack Depth |
|---|---|---|---|---|
| 1 | Reagents | ab_0932 | P8 | 1 |
| 2 | LVLTubes | LVL_MX500_TMT_Tubes | P4 | 1 |
| 3 | Sample | Bio_RadPCR96 | P7 | 1 |

### Final Deck Setup

Below is the final deck layout. Ensure that all the highlighted labware are in the specified positions.

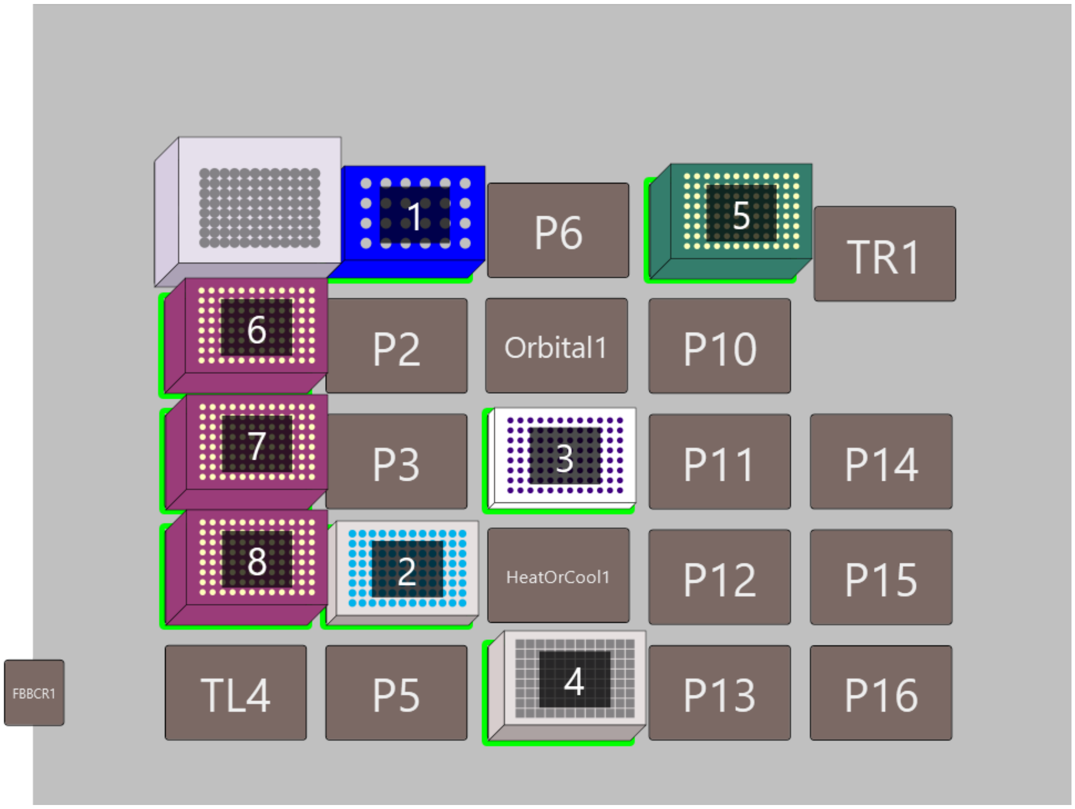


| # | Name | Type | Position | Stack Depth |
|---|---|---|---|---|
| 1 | Final Tubes | BCTubeBlock_2mlTubes | P1 | 1 |
| 2 | LVLTubes | LVL_MX500_TMT_Tubes | P4 | 1 |
| 3 | Sample | Bio_RadPCR96 | P7 | 1 |
| 4 | Reagents | ab_0932 | P8 | 1 |
| 5 | Pooling Tips | BC190F | P9 | 1 |
| 6 | Water Tips | BC50F | TL1 | 1 |
| 7 | TMT Tips | BC50F | TL2 | 1 |
| 8 | AmBi Tips | BC50F | TL3 | 1 |

### Method Details
